# Engineered Chimeric Proteins Functionally Replace ALS-causing Endogenous FUS and TDP-43 to Improve Neuronal Health

**DOI:** 10.64898/2026.09.28.755047

**Authors:** Miyuki Hayashi, Brigid K. Jensen, Amandeep Girdhar, Shashirekha S. Markandaiah, Ying-Hui Ko, Nikhita Arunprakash, Jacquelyn A. DePierro, Tengyang Ni, Wenjing Li, Weixing Zhao, Gino Cingolani, Piera Pasinelli, Lin Guo

## Abstract

Mislocalization and aggregation of the RNA-binding proteins (RBPs) FUS and TDP-43 are pathological hallmarks of amyotrophic lateral sclerosis (ALS) and frontotemporal dementia (FTD), contributing to combined loss- and gain-of-function toxicity. Here, we present a protein engineering strategy that appends optimized anti-aggregation motifs to FUS and TDP-43, thereby enhancing their nuclear localization and solubility, and enabling functional replacement of disease-associated proteins. We show that these engineered chimeric RBPs reduce levels of aggregation-prone endogenous proteins through autoregulatory mechanisms. In addition, chimeric FUS and TDP-43 recapitulate key nuclear functions, including DNA repair and mRNA processing. Notably, chimeric FUS restores neurite outgrowth in diseased iPSC-derived neurons, while chimeric TDP-43 suppresses cryptic exon inclusion in patient-derived fibroblasts. Collectively, these results demonstrate that engineered chimeric RBPs can rebalance RBP homeostasis by simultaneously addressing protein aggregation and mislocalization, thus mitigating RBP-associated loss- and gain-of-function toxicity, highlighting a generalizable strategy for therapeutic protein replacement in neurodegenerative disease.

## INTRODUCTION

Mislocalization and aggregation of RNA-binding proteins (RBPs) are key pathological features of amyotrophic lateral sclerosis (ALS) and frontotemporal dementia (FTD) ^1–14^. Among these, fused in sarcoma (FUS)^14^ and TAR DNA-binding protein 43 (TDP-43)^15^ are central disease drivers, forming cytoplasmic inclusions in affected neurons across both familial and sporadic cases. FUS and TDP-43 share similar domain architectures that enable liquid-liquid phase separation (LLPS) mediated formation of dynamic and functional ribonucleoprotein condensates^16,17^. LLPS allows RBPs to rapidly concentrate locally in response to cellular cues. However, the same features that enable LLPS, including low-complexity domains and multivalent interactions, also render FUS and TDP-43 intrinsically prone to aggregation. When enriched or persistently retained in phase-separated condensates, FUS and TDP-43 can undergo aberrant phase transitions from dynamic liquid assemblies to solid or fibrillar aggregates, thereby inducing toxicity through two non-mutually exclusive mechanisms. First, mislocalization and sequestration into cytoplasmic inclusions deplete functional nuclear proteins, impairing essential processes such as RNA splicing, transcriptional regulation, and genome maintenance and leading to loss-of-function (LoF) toxicity^18–20^. Second, aggregates exert toxic gain-of-function (GoF) effects by aberrantly interacting with other RNAs and proteins. Thus, RBP mislocalization and aggregation can drive neurotoxicity through both LoF and GoF mechanisms, highlighting the need for strategies that address both toxic mechanisms by restoring the diffusive, functional, and nuclear state of RBPs.

A growing body of work, including our own, has demonstrated that nuclear import receptors (NIRs), classically viewed as transport factors, can also function as key regulators of RBP phase transition^21–24^. Coupling this chaperone activity with nuclear transport, NIRs can mitigate toxicity associated with RBP mislocalization and aggregation by simultaneously dissolving their aberrant assemblies and transporting the RBP to the nucleus to restore their nuclear function. For example, the nuclear import receptor Kapβ2 antagonizes FUS aggregation, and increasing Kapβ2 expression restores FUS nuclear localization and function, thereby mitigating its toxicity *in vivo*^24^. Mechanistic studies revealed that NIRs recognize nuclear localization signals (NLSs) not only as transport motifs but also as anti-aggregation signals, and that this NIR-NLS pairing system is highly specific. For example, Kapβ2 selectively recognizes the non-canonical proline-tyrosine NLS (PY-NLS) of FUS but not the canonical NLS (cNLS) of TDP-43, to suppress aggregation by engaging regions that otherwise drive phase separation^25^. Conversely, the cNLS of TDP-43 is recognized by the importin α/β complex (Impα/β), which antagonizes aggregation and promotes nuclear import. Together, these findings establish a general principle: NIR-NLS pairs function as matched chaperone-cargo units that couple nuclear transport with the suppression of aggregation.

Despite its effectiveness under physiological conditions, this coupling between NIR and NLS is compromised by disease-causative mutations or posttranslational modifications that affect the NLS. For example, in ALS, acetylation of lysine 82 within the TDP-43 cNLS disrupts interactions with Impα/β, reducing both nuclear transport and chaperone activity, leading to cytoplasmic mislocalization and aggregation^26,27^. Similarly, severe ALS-causing mutations in FUS either weaken the PY-NLS (e.g., FUS^P525L^) or remove it entirely (e.g., FUS^R495X^)^28^, diminishing Kapβ2 engagement and exacerbating aggregation. Together, these disease-associated perturbations compromise the function of the NLS anti-aggregation signal, creating a gap in the cell’s ability to restrain RBP aggregation.

To address this gap and recover the function of the anti-aggregation signal, we previously conducted a detailed mechanistic investigation to define the rules governing NIR-NLS recognition and anti-aggregation activity^25^. Through this work, we identified two highly efficient NIR-recognition motifs that function as optimized anti-aggregation signals: the 53-amino acid importin-β-binding (IBB) domain, recognized by importin β1 (Impβ1)^29^, and the 38-amino acid synthetic peptide M9M, recognized by Kapβ2^30^. These motifs bind their respective NIRs with higher affinity than native NLSs and engage chaperone activity more effectively. Importantly, tagging RBPs with these signals enables NIRs to chaperone and transport proteins that are not their native cargos. For example, tagging FUS with the IBB domain generates a chimeric RBP FUS_IBB_, which is chaperoned and transported by Impβ1. FUS_IBB_ exhibits improved solubility and nuclear localization compared to the wild-type FUS^25^. Notably, the expression of exogenous FUS reduces the endogenous FUS level^25^, presumably via an autoregulatory feedback loop, a common mechanism for RBPs, including FUS and TDP-43, to maintain proteostasis^31–36^. However, whether this mechanism can be leveraged to engineer soluble RBP chimeras to functionally substitute for endogenous aggregation-prone RBPs, especially in diseased neurons, has not been examined.

In this study, we first optimized and evaluated different FUS_IBB_ constructs in patient-derived iPSC-neurons. Then, we expanded the chimeric protein design to TDP-43, creating and testing TDP-43_M9M_ and TDP-43_IBB_. Our results showed that these engineered RBP chimeras not only downregulate endogenous protein expression but also carry core nuclear functions, such as DNA repair and mRNA regulation. Moreover, these chimeric RBPs restore neurite growth in diseased iPSC-neurons, suggesting that they mitigate toxicities associated with ALS-linked mutations. Together, this work demonstrates that the engineered, highly soluble RBP chimeras can rebalance protein homeostasis in neurons by simultaneously addressing aggregation, mislocalization, and loss-of-function.

## RESULTS

### FUS_500-IBB_ downregulates endogenous FUS in iPSC-derived i^3^ neurons

We previously identified the IBB domain as an effective anti-aggregation signal^25^ and created a highly soluble FUS chimera by replacing the PY-NLS (aa 501-526) with IBB. This chimeric construct, FUS_500-IBB_ (previously called FUS_IBB_), reduced endogenous FUS expression at both the protein and RNA levels in HEK293 and U2OS cells^25^. Before further optimization, we tested whether this original chimeric protein reduces endogenous FUS levels in human neurons. We employed i^3^ cortical neurons (i^3^-CN) generated by stably integrating the human neurogenin-2 transgene under a tetracycline-inducible promoter into patient-derived iPSCs. We transduced i^3^-CN with a lentiviral vector encoding GFP-tagged FUS_500-IBB_ under the human synapsin promoter and assessed FUS expression and localization by immunofluorescence microscopy and Western blotting. Upon transduction, cortical neurons showed FUS_500-IBB_ expression in the nucleus as expected. Moreover, we confirmed a reduction in endogenous FUS protein in FUS_500-IBB_-transduced cells (Figure 1A-D). Thus, FUS_500-IBB_ can effectively downregulate the endogenous WT FUS in iPSC-derived i^3^ cortical neurons.

**Figure 1:**
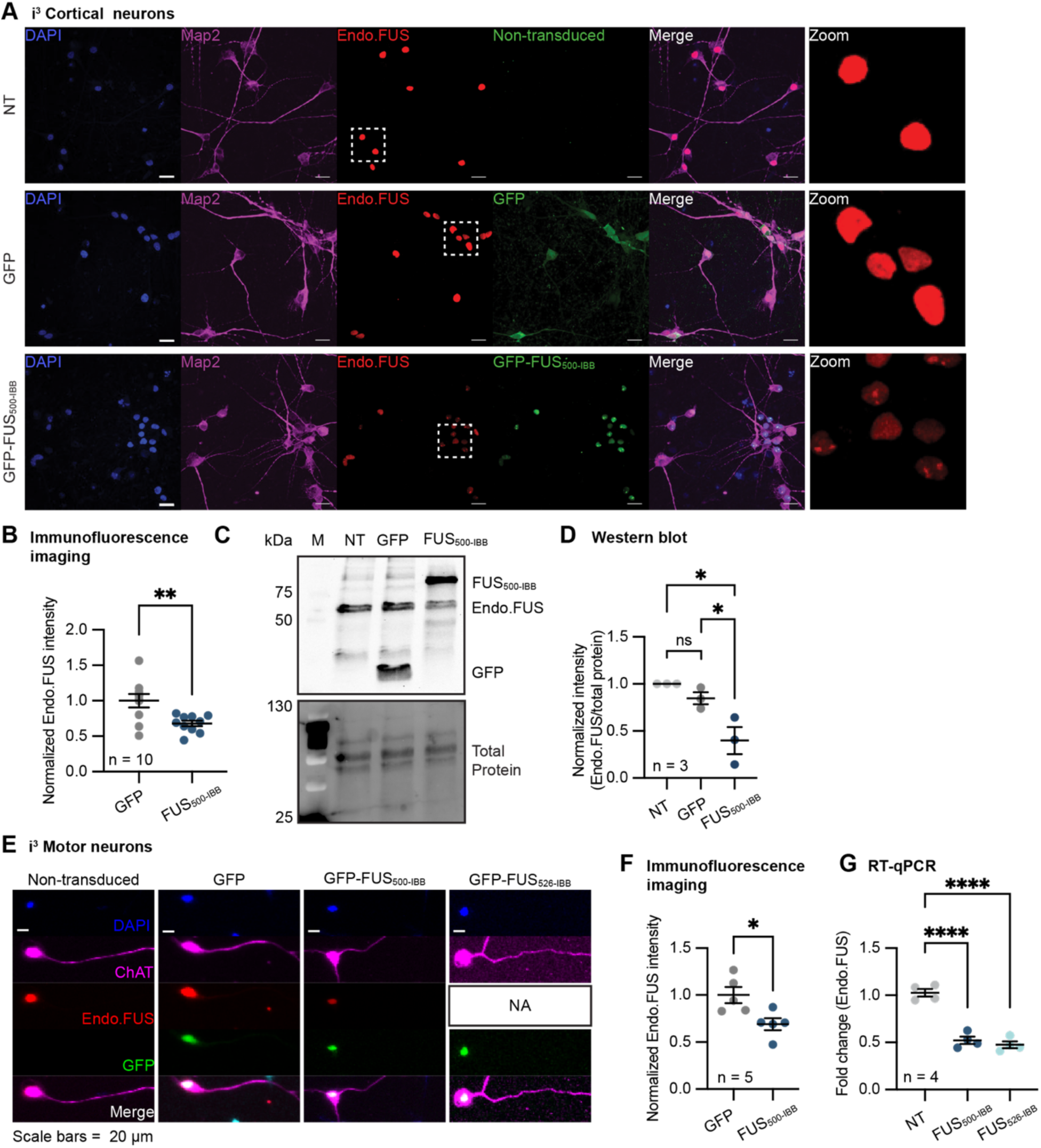
FUS_500-IBB_ and FUS_526-IBB_ downregulate endogenous FUS expression in i^3^ neurons. **A**. Representative immunofluorescence images of i^3^-CN transduced with/without GFP or GFP-FUS_500-IBB_. Boxed area in Endo-FUS panel is enlarged as zoomed image in the final righthand panel. NT = non-transduced control. Scale bars = 20 μm. **B.** Quantification of the endogenous FUS signals shown in A. Mean and standard error of mean (SEM) are plotted, and intensity values are normalized to the mean of GFP-transduced samples. Each point represents the endogenous FUS intensity/GFP area per field, across 3 biological replicates (n = 10 fields total per condition). ** p < 0.01 by Welch’s t test. **C.** Representative Western blot of lysates of i^3^-CN transduced with GFP or FUS_500-IBB_. NT = non-transduced control and M = molecular weight marker. **D.** Quantification of band intensities shown in C. Intensity values were normalized to the total protein. Each point represents an independent experiment (n = 3). * p < 0.05 and ns = not significant by one-way ANOVA Sidak’s multiple comparisons test. NT = non-transduced control. **E.** Representative immunofluorescence images of i^3^-MN transduced with GFP, GFP-FUS_500-IBB_, or GFP-FUS_526-IBB_. Note that the endogenous FUS signal is not shown in FUS_526-IBB_ transduced cells (denoted as NA) because the anti-FUS antibody detects both FUS_526-IBB_ and endogenous FUS. Scale bars = 20 μm. **F.** Quantification of the images collected in E showing endogenous FUS signals from the GFP or GFP-FUS_500-IBB_ transduced neurons. Mean and standard error of mean (SEM) are plotted, and intensity values are normalized to the mean of GFP-transduced samples. Each point represents the average of all fields per well of a condition, with endogenous FUS intensity/GFP area per field measured, across 3 biological replicates (n = 5 wells total per condition, m = 10 fields per well). * p < 0.05 by Welch’s t test. **G.** RT-qPCR analysis of endogenous FUS transcript in i^3^-CN transduced with GFP-FUS_500-IBB_ or GFP-FUS_526-IBB_. Mean and SEM are plotted. Each point represents an independent experiment (n = 4) and Ct values are normalized to GAPDH and non-transfected cells. **** p < 0.0001 by one-way ANOVA Sidak’s multiple comparisons test. NT = non-transduced control. See also **Supplementary Figure 1.**

### Various FUS-IBB chimeras downregulate endogenous FUS levels

FUS_500-IBB_ stays soluble and downregulates the endogenous FUS, exhibiting a potential to be used as a replacement for the aggregation-prone, endogenous FUS. To be used as a replacement, however, the chimeric FUS must function similarly to WT FUS, for example, in RNA regulation^32,37,38^ and DNA repair pathways^39–41^. With only the PY-NLS replaced by IBB and the other domains, including the prion-like domain and the RNA-recognition motif (RRM), staying intact, we anticipate that FUS_500-IBB_ maintains most of FUS function in RNA and DNA regulation. However, several important residues within the PY-NLS, such as arginine 521 (R521), have been suggested to participate in the DNA damage repair process^39^. Thus, we created FUS chimeras that carry additional residues at their C-terminus, making them more akin to WT FUS (Figure S1A). Based on the PY-NLS epitopes^42^, FUS_506-IBB_, FUS_513-IBB_, FUS_522-IBB_, FUS_524-IBB_, and FUS_526-IBB_ were designed (Figure S1A). The expression of all constructs in HEK293 cells was verified by Western blot (Figure S1B). As expected, all constructs are localized to the nucleus (Figure S1C and D). For FUS_506-IBB_ and FUS_513-IBB_, the reduction of the endogenous FUS protein was observed by the immunofluorescence imaging assay (Figure S1C and E), which is comparable to the effect of FUS_500-IBB_. The anti-FUS antibody used in the immunofluorescence study specifically recognizes FUS PY-NLS, enabling the detection of endogenous, full-length FUS distinct from FUS_500-IBB_, FUS_506-IBB_, and FUS_513-IBB_. However, for the constructs containing more PY-NLS residues (i.e., FUS_522-IBB_, FUS_524-IBB_, and FUS_526-IBB_), the antibody can also detect the IBB-tagged exogenous proteins (Figure S1F and G). Thus, immunofluorescence imaging was inadequate for assessing changes in endogenous FUS protein levels in these constructs. Instead, we used Western blot analysis to examine the levels of chimeric and endogenous FUS proteins. To differentiate them from the endogenous FUS by size, the chimeric FUS proteins were tagged with 27 kDa mClover. The expression of mClover-tagged chimeric proteins in HEK293 cells led to the reduction of the endogenous FUS protein (Figure S1F and H). Thus, all FUS chimeras tested here are capable of reducing the endogenous FUS at the protein level.

In the following experiments, to test the activity and functions of FUS chimeras in neuronal cells, we selected the longest construct (i.e., FUS_526-IBB_), which we expect should retain all FUS function, and compared it with the shortest construct (i.e., FUS_500-IBB_) that represents a minimal design. The nuclear localization of both FUS_500-IBB_ and FUS_526-IBB_ was verified in i^3^ motor neurons (i^3^-MN) (Figure 1E). In addition, the autoregulatory activity of FUS_500-IBB_ in i^3^-MN was verified by quantification of the endogenous FUS signals in the immunofluorescence imaging assay (Figure 1E and F). Because of the antibody cross-reactivity of FUS_526-IBB_ discussed above (Figure S1F), the endogenous FUS protein level cannot be assessed via immunofluorescence imaging assay in FUS_526-IBB_ expressing cells. Therefore, we used RT-qPCR with a primer pair specific for endogenous FUS^25^ to test whether FUS_526-IBB_ can regulate endogenous FUS levels. Indeed, the expression of FUS_500-IBB_ and FUS_526-IBB_ reduced the FUS mRNA level compared to the non-transduced control (Figure 1G). In summary, chimeric FUS downregulates the endogenous FUS expression in neuronal cells, including i^3^-CN and i^3^-MN.

### FUS_500-IBB_ and FUS_526-IBB_ downregulate endogenous FUS in disease neurons

FUS^R495X^ and FUS^P525L^ are associated with the most severe forms of ALS. R495X produces a C-terminal truncation, whereas P525L alters the PY-NLS^28^, both of which impair nuclear import and promote cytoplasmic mislocalization and aggregation^43,44^. Therefore, we further tested whether the chimeric protein could downregulate endogenous FUS expression in disease-relevant neurons expressing these mutants. To do so, we obtained iPSC lines carrying FUS^R495X^ and FUS^P525L^, as well as the isogenic Kolf2.1J line, then converted them into i^3^-CN using the transcription factor-mediated method^45^. Upon transduction via lentivirus, the chimeric FUS localized to the nucleus as expected (Figure 2A and B and Figure S2A). Moreover, quantification of the immunofluorescence images showed a 50% decrease in the endogenous FUS level in the FUS_500-IBB_-expressing neurons compared to GFP-expressing neurons (Figure 2C and D). It is worth noting that these lines are heterozygous, and the protein product arising from the FUS^R495X^ allele cannot be detected by the antibody (Figure S2B). Therefore, only the endogenous WT FUS expression can be quantified in FUS^R495X^ neurons using the immunofluorescence assay (Figure 2A and C). Consequently, we quantified the total endogenous FUS expression at the RNA level using RT-qPCR. These RT-qPCR primers are expected to detect both mutant and WT endogenous FUS. In FUS^R495X^ and FUS^P525L^ lines, the chimeric FUS expression led to a 50% downregulation of the endogenous FUS (Figure 2E and F).

**Figure 2:**
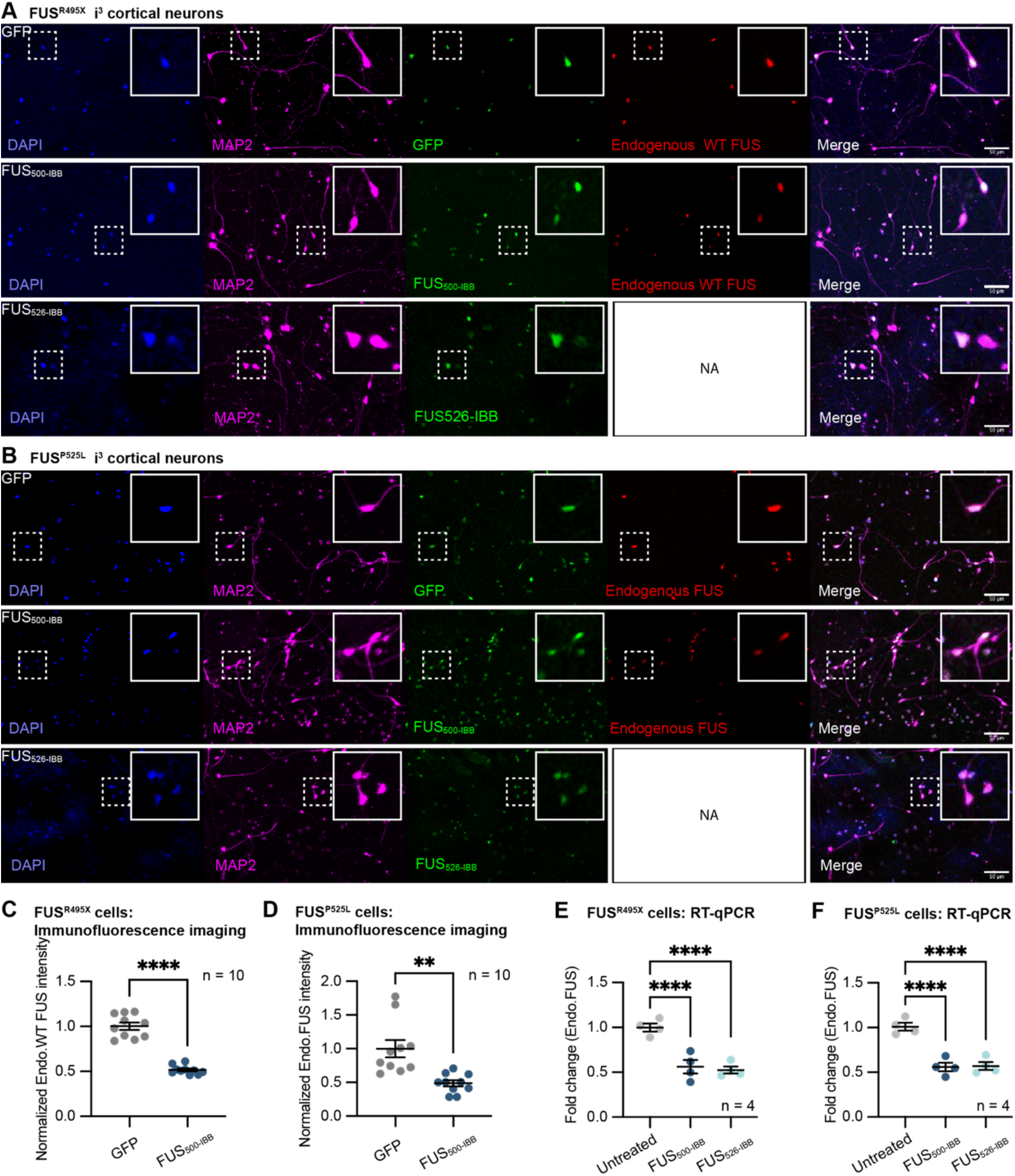
FUS_500-IBB_ and FUS_526-IBB_ downregulate endogenous FUS expression in i^3^-CN carrying FUS^R495X^ and FUS^P525L^ mutation. **A and B.** Representative images of i^3^-CN carrying FUS^R495X^ (A) or FUS^P525L^ (B) transduced with GFP, GFP-FUS_500-IBB_, or GFP-FUS_526-IBB_. Note that the endogenous FUS signal is not shown in FUS_526-IBB_ transduced cells (denoted as NA) because the anti-FUS antibody detects both FUS_526-IBB_ and endogenous FUS. Squares with dotted line indicate the enlarged area. Scale bars = 50 μm. **C.** Quantification of the endogenous WT FUS in FUS^R495X^ i^3^-CN shown in A. Note that the cell line is heterozygous, and the endogenous FUS^R495X^ cannot be detected due to antibody reactivity; thus, the quantification reflects the reduction of FUS from the WT allele. Each point represents the average endogenous WT FUS intensity in transduced cells per field, across 3 biological replicates (n = 10 fields total per condition). **** p < 0.0001 by Welch’s t test. **D.** Quantification of the endogenous FUS in FUS^P525L^ i^3^-CN shown in B. Each point represents the average endogenous FUS intensity in transduced cells per field, across 3 biological replicates (n = 10 fields total per condition). ** p < 0.01 by Welch’s t test. **E and F.** RT-qPCR of the endogenous FUS (including transcripts from both mutant and WT allele) in i^3^-CN harboring FUS^R495X^ (E) or FUS^P525L^ (F) transduced with GFP-FUS_500-IBB_ or GFP-FUS_526-IBB_. Each point represents an independent experiment (n = 4). Ct values are normalized to 18S and Untreated control. ** p < 0.01, *** p < 0.001, and **** p < 0.0001 by one-way ANOVA Sidak’s multiple comparisons test. See also **Supplementary Figure 2**.

### FUS_500-IBB_ and FUS_526-IBB_ function in DNA repair

Having confirmed that FUS_500-IBB_ and FUS_526-IBB_ can downregulate endogenous FUS in both control and disease neurons, we next directly tested whether these chimeric proteins could replace FUS functions. FUS participates in the repair process of double- and single-stranded DNA damages, and impaired strand-break repair is observed in FUS-ALS/FTD models^39,46^. The recruitment of FUS proteins to the DNA break site is a crucial step in initiating repair. Therefore, we tested whether FUS chimeras, including FUS_500-IBB_, FUS_522-IBB_, and FUS_526-IBB_, are recruited to the DNA damage site similarly to WT FUS using a micro-irradiation assay^47^. The FUS_522-IBB_ construct was included because the R521 residue is considered important for the DNA damage repair process^39^. To compare the recruitment of different chimeras to WT FUS and to ensure recruitment is not an overexpression artifact, we prepared stable cell lines using the inducible Flp-In T-Rex HEK293 system that express the respective FUS constructs at endogenous levels upon doxycycline treatment (Figure S3A). To ensure the recruitment of the FUS chimeras to the DNA damage site is an intrinsic property of the chimeras and not dependent on their interaction with endogenous WT FUS, cells were treated with siRNA against FUS (siFUS)^32^, which effectively reduced the endogenous FUS expression without affecting the level of mClover-tagged FUS (Figure S3A-C). After sensitization of the cells with Hoechst, the nuclei were irradiated with a 405-nm laser. As expected, WT FUS was recruited to the damaged site readily after irradiation (Figure 3A-C). Likewise, FUS_500-IBB_, FUS_522-IBB_, and FUS_526-IBB_ were immediately enriched at the damaged site (Figure 3A-B). When the peak recruitment time and the total FUS recruitment were quantified, FUS_522-IBB_ and FUS_526-IBB_ showed similar recruitment kinetics compared to the WT FUS, whereas FUS_500-IBB_ is recruited slightly slower but accumulated longer compared to other FUS constructs tested (Figure 3A-D). Therefore, FUS_522-IBB_ and FUS_526-IBB_ behave more similarly to the WT than FUS_500-IBB_.

**Figure 3:**
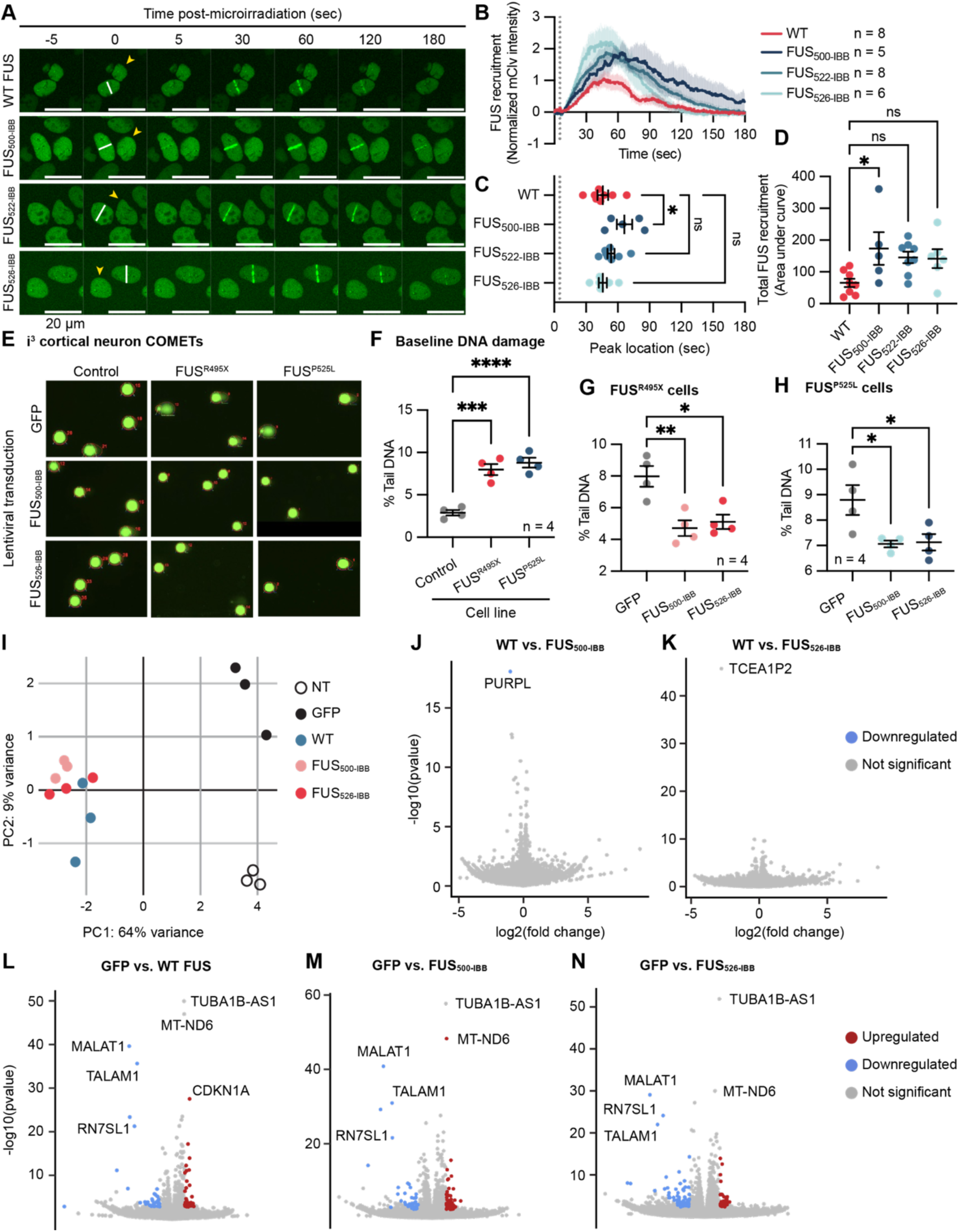
FUS_500-IBB_ and FUS_526-IBB_ participate in DNA repair and regulate mRNA expression similarly to WT FUS. **A.** Representative images from the microirradiation assay. Expression of mClover-tagged FUS was induced by doxycycline in HEK293 cells. At the same time, cells were transfected with siRNA against the endogenous FUS. Upon induction of DNA damage by 405-nm laser, the recruitment of FUS to the irradiated site is recorded over time. The experiment was repeated at least three times with similar results. White line = irradiated site and yellow arrow = reference cell. **B.** Quantification of the images collected in A showing mClover-FUS intensities within the irradiated area over time. Mean and SEM of n = 5∼8 cells are shown. The grey dotted line indicates the time of DNA damage. The experiment was repeated at least three times with similar results. **C.** Quantification of the peak recruitment time shown in B. Each data point represents a cell. The grey dotted line indicates the time of DNA damage. * p < 0.05 and ns = not significant by one-way ANOVA Dunnett’s multiple comparisons test. **D.** Quantification of the area under the curve shown in B. Each data point represents a cell. * p < 0.05 and ns = not significant by one-way ANOVA Sidak’s multiple comparisons test. **E.** Representative images of comet tailing from the COMET assay. Nuclei were harvested from i^3^-CN carrying the indicated FUS mutations transduced with GFP, GFP-FUS_500-IBB_, or GFP-FUS_526-IBB_. **F.** Quantification of E showing the percentage of cells exhibiting tailed DNA in GFP-transduced cells of the indicated cell line. Mean and SEM of four independent experiments are plotted, with approximately 100 nuclei assessed per condition per experiment. *** p < 0.001 and **** p < 0.0001 by one-way ANOVA Dunnett’s multiple comparisons test. **G and H.** Quantification of E showing the percentage of cells exhibiting tailed DNA in GFP, GFP-FUS_500-IBB_, or GFP-FUS_526-IBB_-transduced i^3^-CN harboring FUS^R495X^ (G), or FUS^P525L^ (H). Mean and SEM of four independent experiments are plotted with approximately 100 nuclei assessed per condition per experiment. * p < 0.05 and ** p < 0.01 by one-way ANOVA Dunnett’s multiple comparisons test. **I.** Principal component analysis based on differentially expressed genes in samples. **J and K.** Differential expression analysis of the indicated FUS chimeras compared to WT FUS. Data points are colored based on log_2_(fold change) > 1 and significance (p < 0.05). **L-N.** Differential expression analysis of WT FUS and the indicated FUS chimeras compared to GFP transduced cells. Data points are colored based on log_2_(fold change) > 1 and significance (p < 0.05). See also **Supplementary Figure 3**.

Next, we tested whether recruitment of the FUS chimeras influences repair of damaged DNA using a GFP-based homologous recombination (HR) reporter assay^48,49^. In this system, a double-strand break (DSB) is generated at an engineered I-SceI restriction site within a nonfunctional GFP cassette stably integrated in U2OS cells (Figure S3D). HR-mediated repair restores an intact GFP expression cassette, and the fraction of GFP-positive cells is quantified by flow cytometry. siRNA treatment to knock down FUS results in a 67% reduction in DSB repair efficiency, confirming the role played by FUS in DNA repair (Figure S3E). Notably, expression of either FUS_500-IBB_ or FUS_526-IBB_ restored HR to a level comparable to WT FUS (Figure S3E), suggesting the modest difference in the recruitment kinetics observed for FUS_500-IBB_ does not compromise its ability to support HR repair.

Finally, we asked whether FUS chimeras can rescue impaired DNA repair caused by FUS mutations (FUS^R495X^ and FUS^P525L^) in neuronal cells. In i^3^-CN carrying FUS mutations, the elevated baseline DNA damage was evidenced by an increased number of nuclei exhibiting comet tailing compared to the control (Figure 3E and F). Expression of chimeric FUS significantly reduced DNA damage levels compared with mutant FUS lines expressing GFP alone (Figure 3E, G, and H). In summary, FUS_500-IBB_ and FUS_526-IBB_ restore the function of WT FUS in DNA repair in disease-relevant neurons.

### FUS_500-IBB_ and FUS_526-IBB_ regulate mRNA expression like WT FUS

FUS regulates the levels of transcripts through direct binding^31,50^ or indirectly by affecting microRNA stability^51^. Given that full-length RRMs are present in the constructs, we hypothesized that FUS-IBB chimeras could regulate the transcriptome similarly to WT FUS. To test this, we extracted RNA from i^3^-CN expressing GFP (Figure S3F), WT FUS, FUS_500-IBB_, or FUS_526-IBB_ and subjected them to RNA sequencing. In the principal component analysis, reads from neurons expressing FUS clustered together, distinct from those of non-transduced (NT) and GFP-expressing neurons (Figure 3I). Because the lentiviral transduction itself altered the mRNA expression (Figure S3G), we chose to use either WT FUS or GFP-expressing neurons as a control for the chimeric FUS to perform differential expression analysis. The differential expression analysis revealed one transcript, *purpl*, downregulated in FUS_500-IBB,_ and no significant difference in FUS_526-IBB_ compared to the WT FUS-expressing neurons (Figure 3J and K). When comparing to the GFP-expressing neurons, all FUS-expressing neurons showed a similar transcriptomic landscape, namely the upregulation of *mt-nd6* and *cdkn1a*, as well as the downregulation of *malat1*, *talam1*, and *rn7sl1* (Figure 3L-N). We also compared the differential expression of selected transcripts to the previously characterized FUS regulatory targets. *Caskin1*, *cttn*, *ptbp1*, *tnfsf14*, and *nlrp2* showed trends in upregulation, and *sh3kbp1*, *tia1*, and *s1pr1* showed trends in downregulation, consistent with previous reports (Figure S3H)^50,52,53^. Therefore, the chimeric FUS_500-IBB_ and FUS_526-IBB_ regulate mRNA similarly to the WT FUS.

### FUS_500-IBB_ and FUS_526-IBB_ expression restores neurite growth

FUS_500-IBB_ and FUS_526-IBB_ can downregulate endogenous FUS in disease neurons and function similarly to WT FUS in DNA repair and mRNA regulation. We therefore tested whether the expression of the chimeric FUS could restore neuronal health. Longitudinal survival analysis shows that expression of chimeric FUS did not significantly alter their viability, supporting the safety of FUS chimera treatment in neurons (Figure S4). Furthermore, we assessed overall neurite lengths in i^3^-CN bearing FUS mutations as a morphological indicator of neuronal health. At baseline, where GFP protein was overexpressed, transduced FUS^R495X^ and FUS^P525L^ neurons had shorter neurites than control neurons, suggesting potential toxicity from the mutant (Figure 4A-D). Notably, overexpression of WT FUS reduced the average neurite length in both control and mutant neurons (Figures 4A-C and E-G), which is consistent with previous observations that overexpression of WT FUS can be toxic^54^. On the contrary, chimeric FUS-expressing neurons did not show neurite growth impairment in control neurons (Figures 4A and E), confirming its safety. Remarkably, in disease neurons expressing FUS mutant FUS^R495X^ or FUS^P525L^, expression of either chimeric FUS rescued the neurite length comparable to that of the control neurons (Figure 4A-C and E-G). Taken together, the chimeric FUS_500-IBB_ and FUS_526-IBB_ downregulate the expression of mutant FUS, replace FUS function, and restore neuronal health and viability.

**Figure 4:**
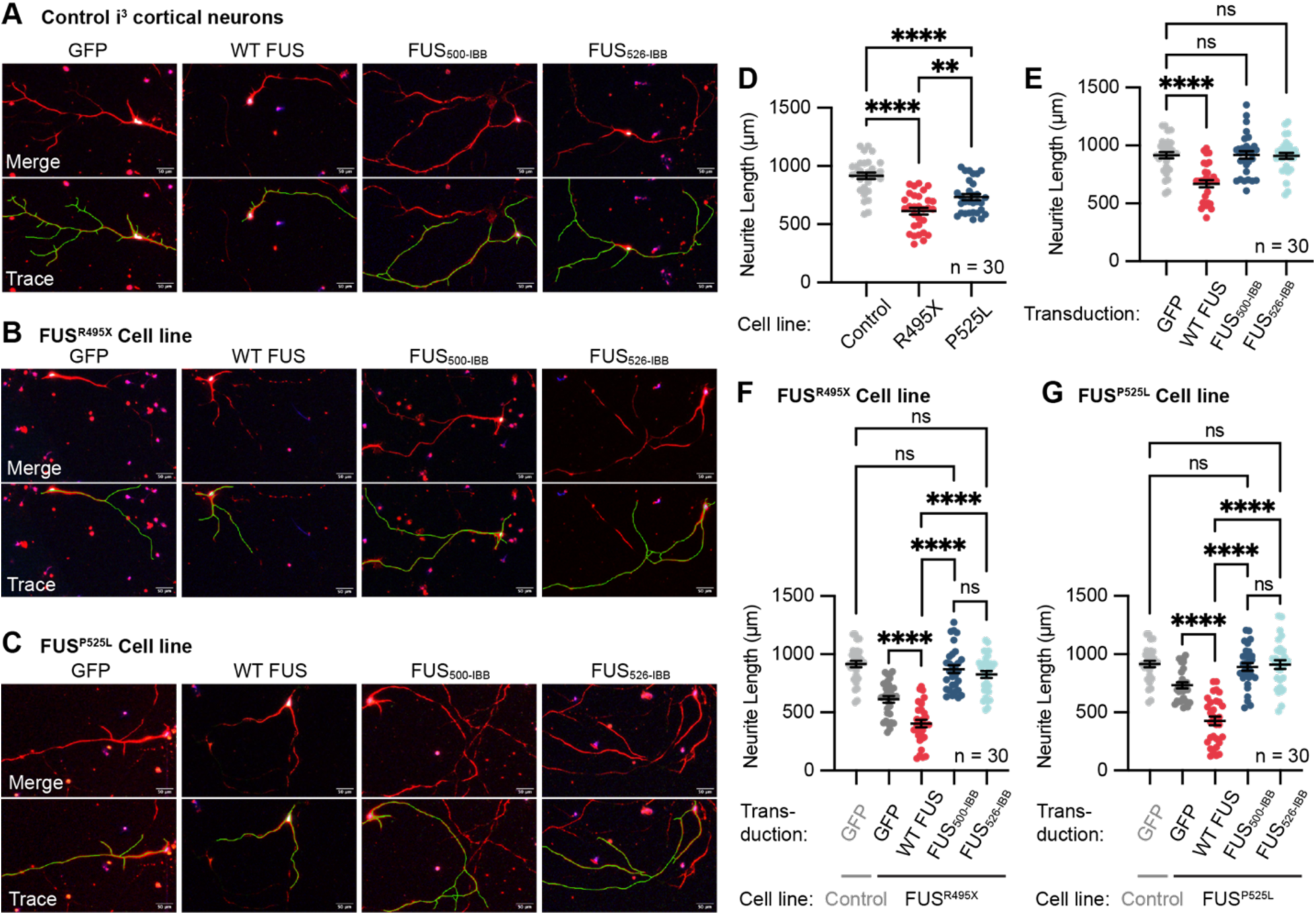
Chimeric FUS expression is not toxic and restores neurite growth of i^3^-CN harboring mutant FUS. **A-C.** Representative immunofluorescence images of control (A), FUS^R495X^ (B), and FUS^P525L^ (C) i^3^-CN transduced with GFP-tag control, GFP-tagged WT FUS, or chimeric FUS. Top panels represent merged channel images (blue = DAPI, red = MAP2, and green = GFP-tagged FUS). The bottom panels include neurite tracing (green line) over the merged channel images. Scale bars = 50 μm. **D.** Baseline neurite length quantification. Neurite lengths were measured as represented in A-C for control and FUS mutant lines expressing GFP. Each point represents a transduced neuron. Data were collected across 3 independent experiments, 10 cells per experiment (biological replicates), n = 30 cells total per condition. Mean and SEM are plotted. ** p < 0.01 and **** p < 0.0001 by one-way ANOVA Tukey’s multiple comparisons test. **E.** Quantification of images in A showing neurite length in control neurons transduced with GFP, or GFP-tagged WT, or chimeric FUS. Each point represents a transduced neuron. Data were collected across 3 independent experiments, 10 cells per experiment (biological replicates), n = 30 cells total per condition. Mean and SEM are plotted. ns = not significant and **** p < 0.0001 by one-way ANOVA Dunnett’s multiple comparisons test. **F and G.** Quantification of neurite lengths shown in A and B (F), and A and C (G). Each point represents a transduced neuron. Data was collected across 3 independent experiments, 10 cells per experiment (biological replicates), n = 30 cells total per condition. Mean and SEM are plotted. **** p < 0.0001 by one-way ANOVA Sidak’s multiple comparisons test. See also **Supplementary Figure 4.**

### TDP-43_M9M_ and TDP-43_IBB_ are chaperoned by nuclear import receptors, localize to the nucleus, and reduce endogenous TDP-43 expression

Given the success of engineered FUS chimeras in restoring solubility, nuclear function, and neuronal health, we next asked whether a similar NIR-NLS-based strategy could be generalized beyond FUS and extended to TDP-43. TDP-43 aggregation underlies the predominant pathology observed in ALS and is similar to FUS in its domain architecture and autoregulated expression^55^. We have demonstrated that the aggregation of a TDP-43 chimera, TDP-43_M9M_, can be prevented and reversed by Kapβ2 in a purified protein system^25^. We envision that, as with FUS, highly soluble chimeric TDP-43 proteins can be used to downregulate endogenous TDP-43 expression and replace its function. To broaden the design space and compare distinct NIR-recognition strategies, we additionally generated a second TDP-43 chimera bearing the importin-β-binding domain (IBB), termed TDP-43_IBB_ (Figure 5A). We first purified TDP-43_IBB_-MBP and tested its aggregation properties *in vitro*. After cleaving the solubility tag MBP, TDP-43_IBB_ formed aggregates, as quantified by the increase in turbidity (Figure 5B). Addition of Impβ1 prevented the formation of TDP-43_IBB_ aggregates but not WT TDP-43 (Figure 5B-D and S5A and B), which is similar to the activity of Kapβ2 on TDP-43_M9M_ characterized previously^25^. Somewhat unexpectedly, however, Impβ1 failed to disaggregate the pre-formed TDP-43_IBB_ aggregates (Figure 5E and F), unlike the TDP-43_M9M_-Kapβ2 pair^25^. Therefore, at equal molar concentrations, the IBB-Impβ1 pair can prevent, but cannot reverse, TDP-43 aggregation *in vitro*.

**Figure 5:**
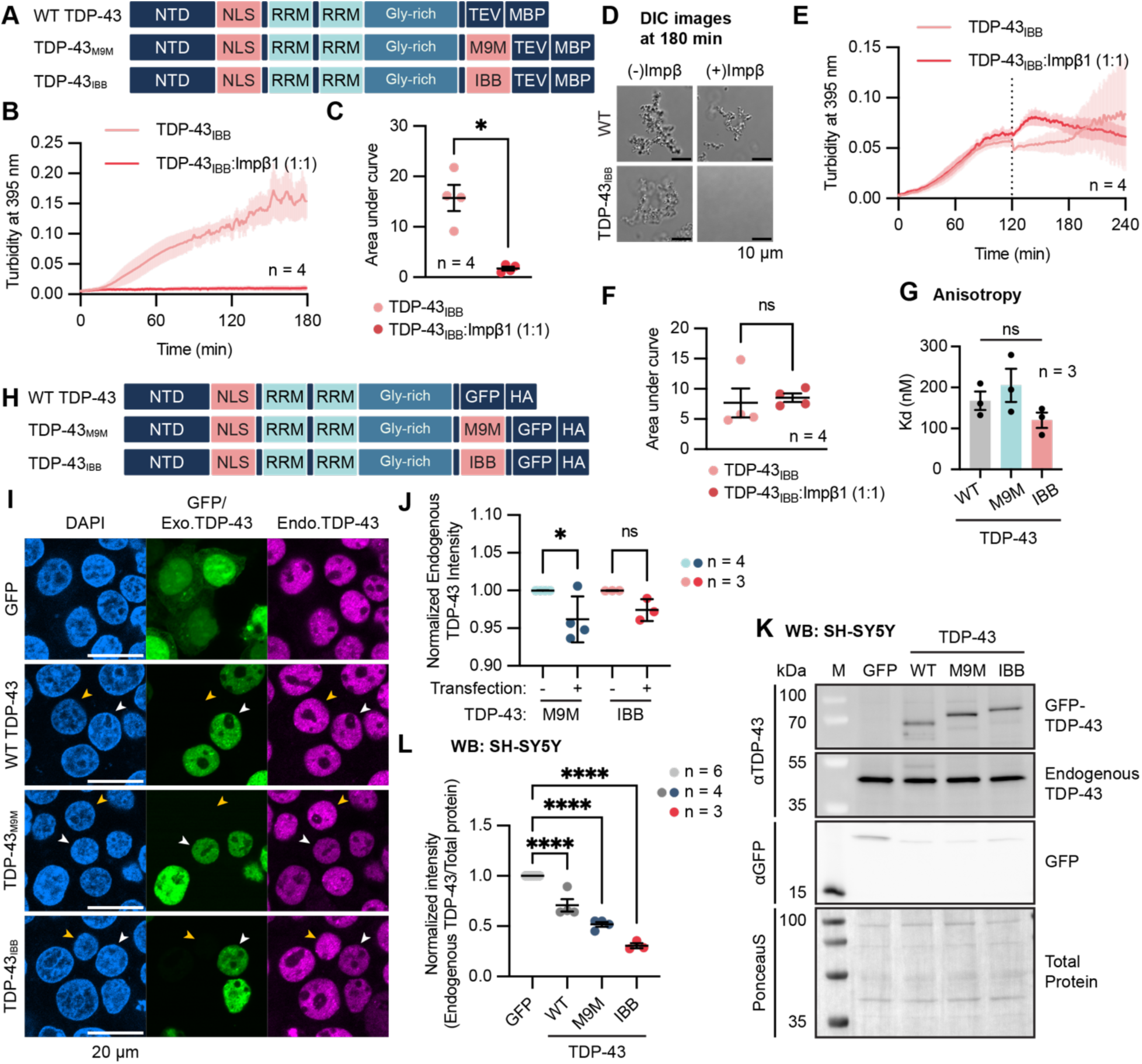
TDP-43_M9M_ and TDP-43_IBB_ reduce endogenous TDP-43 expression. **A.** Design of M9M- and IBB-tagged TDP-43 constructs for bacterial expression and purification. Constructs are tagged with MBP at the C-terminus, and the cleavage of the tag by TEV protease triggers TDP-43 phase separation and aggregation. NTD = N-terminal domain, NLS = nuclear localization signal, RRM = RNA recognition motif, Gly-rich = glycine-rich low complexity domain, and TEV = TEV protease target site. Domains are not scaled to the number of amino acids. **B.** Turbidity assay of TDP-43_IBB_ assembly with and without an equimolar Impβ1. TDP-43_IBB_ and Impβ1 were mixed at the beginning of the reaction when TEV was added to initiate TDP-43 assembly. Turbidity was recorded over time. Mean and SEM of 4 independent experiments are plotted. **C.** Quantification of the turbidity curves in B showing the area under the curve calculation of the indicated condition. Each point represents an independent experiment. * p < 0.05 by Welch’s t test. **D.** Representative DIC images of TDP-43 aggregates taken at the end of the turbidity assay (at 180 min). Scale bars = 10 μm. **E.** Turbidity assay curve of TDP-43_IBB_. TDP-43_IBB_ aggregates were assembled for 120 min before the addition of equimolar Impβ1 (dotted line), and changes in turbidity were measured over time. Mean and SEM of 4 independent experiments are plotted. **F.** Quantification of the area under the curve shown in E. Each point represents an independent experiment. ns = not significant by Welch’s t test. **G.** Quantification of anisotropy assays in Supplementary Figure 5C showing the Dissociation constant (Kd) of CLIP-34 binding to WT or indicated chimeric TDP-43. Each data point represents an independent experiment. ns = not significant by one-way ANOVA Tukey’s multiple comparisons test. **H.** Design of M9M- and IBB-tagged TDP-43 for mammalian expression. HA = hemagglutinin tag. Domains are not scaled to the number of amino acids. **I.** Representative fluorescence images of HEK293T cells with endogenously Halo-tagged TDP-43 transfected with GFP or the indicated GFP-tagged TDP-43 constructs. Cells were fixed, and endogenous TDP-43 was visualized by the addition of Halo-ligand conjugated with Janelia Fluor 646. White arrows = transfected cells and yellow arrows = non-transfected cells. Scale bars = 20 μm. **J.** Quantification of the endogenous TDP-43 fluorescence signal shown in I. Each point represents an independent experiment. Mean pixel intensity values are normalized to that of the non-transfected cells. * p < 0.05 and ns = not significant by one-way ANOVA Sidak’s multiple comparisons test. **K.** Representative Western blot of SH-SY5Y cells expressing GFP or the indicated GFP-tagged TDP-43 constructs. M = molecular weight marker. Experiments were repeated at least three times with similar results. **L.** Quantification of the Western blots shown in K. Each point represents an independent experiment. **** p < 0.0001 by one-way ANOVA Tukey’s multiple comparisons test. See also **Supplementary Figure 5.**

Next, we tested whether TDP-43 chimeras can engage the autoregulation feedback loop to downregulate endogenous TDP-43 levels. During autoregulation, recruitment of TDP-43 to its transcript is mediated by CLIP-34 RNA encoded at the 3’ UTR of the TDP-43 mRNA^36,56^. Using an anisotropy assay, we confirmed that both TDP-43 chimeras, TDP-43_IBB_ and TDP-43_M9M_, exhibited binding affinity for CLIP-34 RNA similar to that of WT TDP-43 (Figure 5G and S5C), supporting their ability to engage the autoregulation mechanism. To further test their cellular localization and ability to regulate endogenous TDP-43 levels, we overexpressed the chimeric TDP-43 proteins in HEK293T cells. Unlike the FUS antibody used in the experiments above, the TDP-43 antibody cannot distinguish endogenous from exogenous TDP-43. Thus, we utilized a HEK293T cell line, in which endogenous TDP-43 is Halo-tagged^57^. Upon transfection, GFP-tagged chimeric TDP-43 (Figure 5H) localized to the nucleus (Figure 5I). Quantification of the fluorescence intensity of Halo-tagged endogenous TDP-43 shows that TDP-43_M9M_ significantly reduced endogenous TDP-43 expression (Figure 5J). TDP-43_IBB_ also trended toward reduced endogenous TDP-43 expression, although the effect did not reach statistical significance (Figure 5J). It is unlikely but possible that the Halo-tag may have affected the binding of the chimeric TDP-43 to the endogenous TDP-43 transcript, thereby altering autoregulation. Thus, we further tested the chimeric proteins in SH-SY5Y neuroblastoma cells that carry untagged TDP-43. Notably, both chimeras and WT TDP-43 reduced the endogenous TDP-43 expression (Figure 5K-L). In summary, the chimeric TDP-43_M9M_ and TDP-43_IBB_ reduce endogenous TDP-43 levels.

The M9M peptide is an efficient anti-aggregation signal with high affinity to Kapβ2^25^. However, it must not overwhelm the Kapβ2-mediated transport pathway to disturb the nuclear-cytoplasmic localization of other Kapβ2 cargos. Thus, we monitored the localization of a native Kapβ2 cargo, FUS^42^, to ensure that FUS localization is not affected by the expression of a competing cargo, TDP-43_M9M_. In TDP-43_M9M_ expressing cells, FUS nuclear intensity or nuclear-to-cytoplasmic ratio does not decrease compared to the non-transfected cells (Figure S5D-F). Therefore, TDP-43_M9M_ does not reduce the nuclear localization of a native Kapβ2 cargo.

### TDP-43 chimeras maintain nuclear localization of themselves and endogenous TDP-43 under proteasomal inhibition

Age-dependent decline in proteasome activity has been implicated as a trigger of TDP-43 proteinopathy, driving nuclear LoF and cytoplasmic accumulation^26^. Reduced proteasomal activity disrupts Impα/β-mediated nuclear import of TDP-43 by altering post-translational modifications, specifically lysine acetylation within its NLS^26,27^. Notably, this mechanism has been proposed as a potential initiator of TDP-43 pathology in sporadic ALS cases lacking mutations in TDP-43 itself^26^. Therefore, we tested whether engineered TDP-43 chimeras confer resistance to the mislocalization and aggregation caused by reduced proteasomal activity. To model this aging-associated cellular context that compromises endogenous cNLS function, we treated cells with the proteasome inhibitor bortezomib (BTZ^26^. Consistent with the previous report, BTZ treatment caused WT TDP-43 to mislocalize to the cytoplasm (Figure 6A). In contrast, both TDP-43_M9M_ and TDP-43_IBB_ retained predominantly nuclear localization under the same conditions (Figure 6A-C). Notably, endogenous TDP-43 also exhibited an increased nuclear presence in the chimeric TDP-43-expressing cells, indicating that the expression of chimeric TDP-43 can promote the nuclear localization of endogenous TDP-43 (Figure 6A, D, and E). Together, these results indicate that TDP-43 chimeras resist mislocalization of both the engineered protein and endogenous TDP-43 in an aging-relevant cellular context characterized by impaired proteasomal activity.

**Figure 6:**
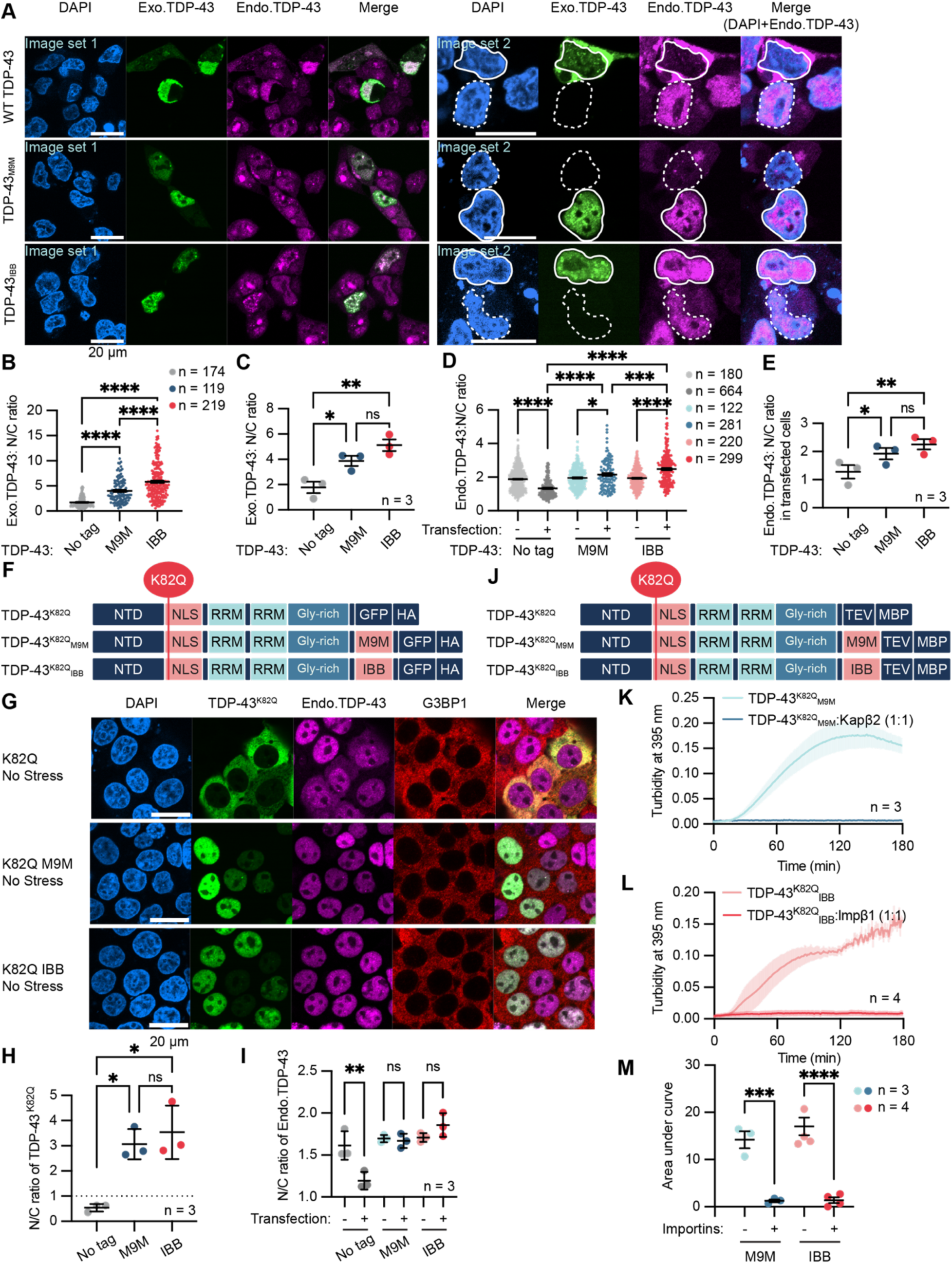
TDP-43_M9M_ and TDP-43_IBB_ resist proteasomal inhibition-induced mislocalization. **A.** Representative fluorescence images of HEK293T carrying endogenously Halo-tagged TDP-43 transfected with the indicated TDP-43 constructs. Larger field view (left; image set 1) and zoomed view (right; image set 2). In image set 2, nuclei of transfected cells are circled by solid lines, and non-transfected cells are circled by dashed lines. Scale bars = 20 μm. **B.** Quantification of the exogenous TDP-43 (Exo.TDP-43) nuclear-to-cytoplasmic (N/C) ratio from a representative experiment as shown in A. Each data point represents a cell. **** p < 0.0001 by one-way ANOVA Tukey’s multiple comparisons test. **C.** Quantification of Exo. TDP-43 N/C ratio from three independent experiments. ns = not significant, * p < 0.05, and ** p < 0.01 by one-way ANOVA Tukey’s multiple comparisons test. **D.** Quantification of the endogenous TDP-43 (Endo.TDP-43) N/C ratio from a representative experiment as shown in A. Each data point represents a cell. *p < 0.05 and **** p < 0.0001 by one-way ANOVA Sidak’s multiple comparisons test. **E.** Quantification of Endo.TDP-43 N/C ratio from three independent experiments. ns = not significant, * p < 0.05, and ** p < 0.01 by one-way ANOVA Sidak’s multiple comparisons test. **F.** Design of TDP-43^K82Q^ constructs. Lysine 82 within the NLS is mutated to glutamine. HA = hemagglutinin tag. Domains shown are not scaled to the number of amino acids. **G.** Representative immunofluorescence images of HEK293T carrying endogenously Halo-tagged TDP-43 transfected with the indicated TDP-43^K82Q^ constructs with/without IBB or M9M tag. G3BP1 = stress granule marker. **H.** Quantification of images collected in G showing N/C ratio of TDP-43^K82Q^ constructs with/without IBB or M9M tag. Mean and SEM are plotted. Each point represents an independent experiment. ns = not significant and * p < 0.05 by one-way ANOVA Tukey’s multiple comparisons test. **I.** Quantification of images collected in G showing N/C ratio of endogenous TDP-43 in TDP-43^K82Q^ transfected and non-stressed cells. Each point represents an independent experiment. ns = not significant and ** p < 0.01 by one-way ANOVA Sidak’s multiple comparisons test. **J.** Design of TDP-43^K82Q^ constructs for bacterial expression. Domains shown are not scaled to the number of amino acids. **K.** Turbidity assay of TDP-43^K82Q^_M9M_ with and without an equimolar amount of Kapβ2. TDP-43^K82Q^_M9M_ and Kapβ2 were mixed at the beginning of the reaction, when TEV was added to initiate TDP-43 assembly. Turbidity was recorded over time. Mean and SEM of three independent experiments are plotted. **L.** Turbidity assay of TDP-43^K82Q^_IBB_ with and without equimolar amount of Impβ1. TDP-43^K82Q^_IBB_ and Impβ1 were mixed at the beginning of the reaction, when TEV was added to initiate TDP-43 assembly. Turbidity was recorded over time. Mean and SEM of four independent experiments are plotted. **M.** The area under the curve quantification of graphs shown in K and L. Each point represents an independent experiment. *** p < 0.001 and **** p < 0.0001 by one-way ANOVA Sidak’s multiple comparisons test. See also **Supplementary Figure 6.**

### TDP-43 chimeras resist K82Q-induced mislocalization and aggregation

While BTZ treatment models a physiologically relevant aging-associated perturbation that enhances TDP-43 acetylation, reduced proteasomal activity affects multiple cellular pathways. To isolate the contribution of TDP-43 acetylation itself, we next focused on acetylation of K82 within the cNLS, which has been identified as the most disruptive modification for Impα/β binding and nuclear localization^26,27^. We therefore directly tested whether engineered TDP-43 chimeras could resist the mislocalization- and aggregation-promoting effects of the disease-relevant acetyl-mimetic mutation K82Q^26,27^.

We expressed TDP-43_M9M_ or TDP-43_IBB_ that harbor the K82Q mutation, namely TDP-43^K82Q^_M9M_ and TDP-43^K82Q^_IBB_ (Figure 6F), in HEK293T cells carrying endogenously Halo-tagged TDP-43. Consistent with prior reports, TDP-43^K82Q^ localized predominantly to the cytoplasm (Figure 6G). In contrast, both TDP-43^K82Q^_M9M_ and TDP-43^K82Q^_IBB_ localized efficiently to the nucleus (Figure 6G and H). Even under stressed conditions, TDP-43^K82Q^_M9M_ and TDP-43^K82Q^_IBB_ maintain nuclear localization and are able to condense into nuclear stress bodies, whereas TDP-43^K82Q^ forms cytoplasmic puncta adjacent to the stress granules (Figure S6). Notably, the expression of TDP-43^K82Q^ led to the mislocalization of the endogenous TDP-43, which was not observed in TDP-43^K82Q^_M9M_ or TDP-43^K82Q^_IBB_-expressing cells (Figure 6G and I).

To determine whether the resistance to mislocalization reflected direct chaperone activity, we examined aggregation of TDP-43^K82Q^ chimeras in a pure protein assay (Figure 6J-M). Similar to non-K82Q constructs, equimolar amounts of the corresponding nuclear import receptors prevented aggregation of TDP-43^K82Q^_M9M_ and TDP-43^K82Q^_IBB_ (Figure 6K-M). These findings demonstrate that optimized NIR-NLS pairs enable NIR to chaperone and transport TDP-43 even when the native cNLS is functionally compromised by acetylation.

### TDP-43_M9M_ reduces cryptic exon-containing transcripts in patient-derived fibroblasts

Given that chimeric TDP-43 proteins can (1) be chaperoned and transported to the nucleus by NIRs, and (2) reduce endogenous TDP-43 expression, they may be used as a therapeutic agent to functionally replace endogenous TDP-43. Thus, the chimeric TDP-43 must not exhibit toxicity when expressed in neurons. To test this, we transduced tdTomato-tagged WT and chimeric TDP-43 in i^3^-CN and assessed whether neurite growth was affected by chimeric protein expression. In WT TDP-43-expressing neurons, neurite length was significantly reduced compared to tdTomato control-expressing neurons (Figure 7A and B and Supplementary Figure 7A), consistent with previous observations that overexpression of WT TDP-43 confers neuronal toxicity^58^. In contrast, neurons expressing chimeric TDP-43 exhibited neurite lengths comparable to those of tdTomato control-expressing neurons (Figure 7A and B and Supplementary Figure 7A). Therefore, the expression of TDP-43_M9M_ nor TDP-43_IBB_ is toxic to neurons.

**Figure 7:**
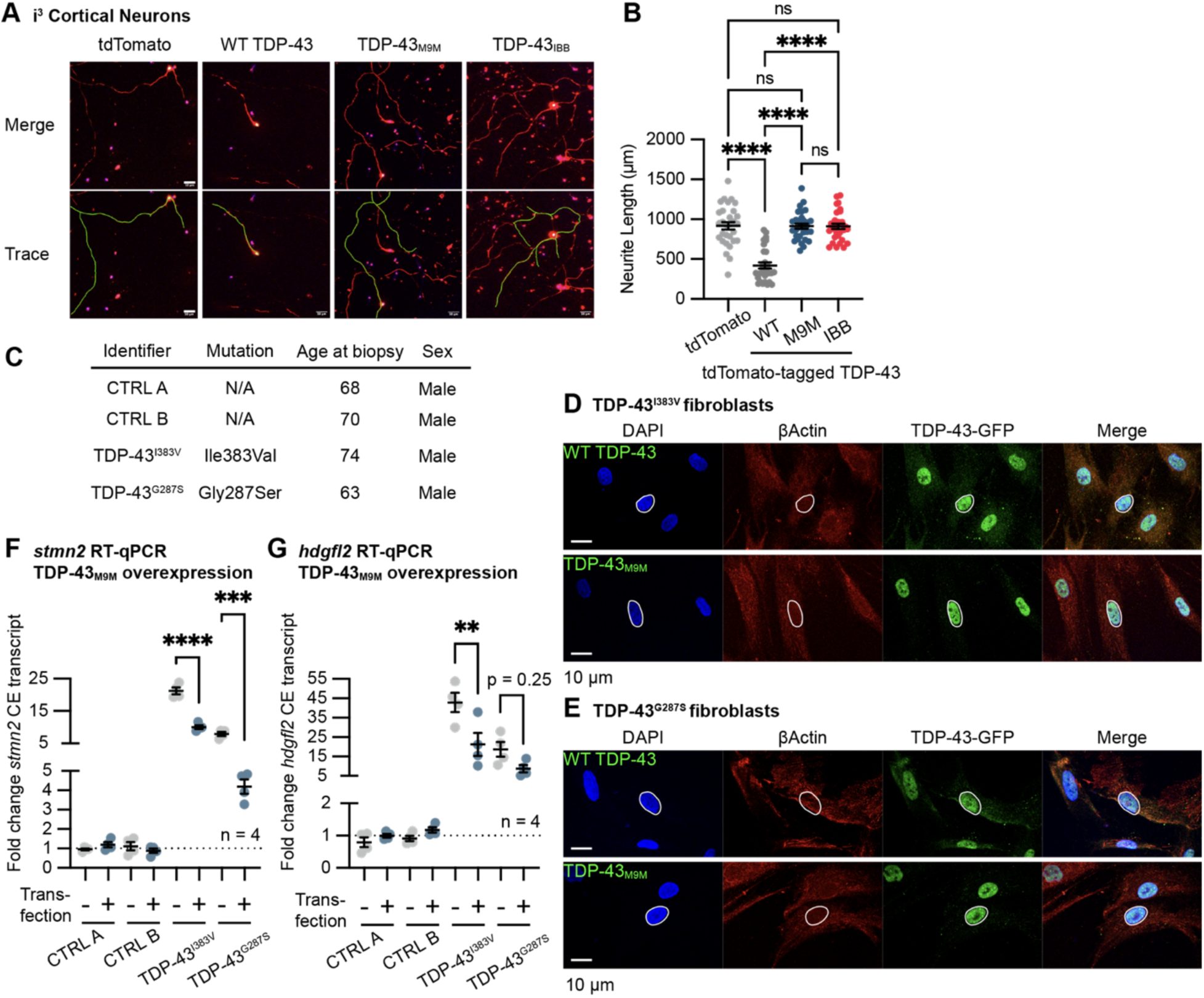
Chimeric TDP-43 are not toxic to neurons and can regulate cryptic exon splicing. **A.** Representative immunofluorescence images of control i^3^-CN transduced with tdTomato-tagged WT or chimeric TDP-43. Top panels represent merged channel images (blue = DAPI, red = MAP2, and green = tdTomato-tagged TDP-43), and the bottom panels represent examples of traced neurites (green line). Scale bars = 50 μm. **B.** Quantification of neurite lengths. Neurite lengths were measured as represented in A. Each point represents a transduced neuron. Data were collected across 3 independent experiments, 10 cells per experiment, n = 30 cells total per condition. Mean and SEM are plotted. ns = not significant and **** p < 0.0001 by one-way ANOVA Tukey’s multiple comparisons test. **C.** Demographics of patient-derived fibroblasts used to test TDP-43_M9M_ activity. Two TDP-43 mutation carriers were matched in age and sex with non-ALS control individuals. **D and E.** Representative immunofluorescence images of ALS-patient fibroblasts, carrying TDP-43^I383V^ (D) and TDP-43^G287S^ (E) mutations, transfected with GFP-tagged WT or chimeric TDP-43. The nucleus of a representative cell is circled with a white line. Scale bars = 10 μm. **F and G.** Quantification of *stmn2* (F) and *hdgfl2* (G) cryptic exon-containing transcripts. ALS-patient fibroblasts were transfected with TDP-43_M9M_, and RNAs were extracted for the quantification by RT-qPCR. Each point represents an independent experiment (n = 4). Ct values were normalized to GAPDH and CTRLs A and B without transfection. ** p < 0.01, *** p < 0.001, and **** p < 0.0001 by one-way ANOVA Sidak’s multiple comparisons test. See also **Supplementary** Figure 7.

In the purified protein system, aggregation of TDP-43_M9M_ can be prevented and reversed by an equimolar amount of Kapβ2^25^. Moreover, TDP-43_M9M_ more robustly reduced endogenous TDP-43 level across different cell lines (Figure 5I-M). Given the superior performance of the TDP-43_M9M_, we decided to take it further to test if TDP-43_M9M_ can function like WT TDP-43. TDP-43 regulates pre-mRNA splicing, thereby reducing the inclusion of cryptic exons in transcripts such as *stmn2* and *hdgfl2*^18,20,59,60^. We obtained two fibroblast lines from ALS patients carrying the I383V or G287S mutations, and two age- and sex-matched controls, to test the ability of TDP-43_M9M_ to regulate cryptic exon splicing (Figure 7C). As expected, TDP-43_M9M_ localized to the nucleus of the ALS-patient and control fibroblasts (Figure 7D and E and Supplementary Figure 7B and C). In fibroblasts carrying TDP-43 mutations, an increased level of cryptic exon inclusion in *stmn2* and *hdgfl2* transcripts was observed (Figure 7F and G). With the expression of TDP-43_M9M_, however, the cryptic exon inclusion in *stmn2* transcripts was significantly reduced (Figure 7F). In *hdgfl2* transcripts, the effect of TDP-43_M9M_ expression differed between the mutant lines, with TDP-43^I383V^ fibroblasts showing a significant reduction, whereas TDP-43^G287S^ showed a downward but not significant reduction (Figure 7G). In summary, TDP-43_M9M_ rescues the loss of the endogenous TDP-43 function in fibroblasts carrying mutant TDP-43.

## DISCUSSION

In this study, we present chimeric proteins of FUS and TDP-43 tagged with highly efficient anti-aggregation signal, namely FUS_IBB_, TDP-43_M9M_, and TDP-43_IBB_, that can be engaged and chaperoned by their recognition NIRs even under diseased conditions. Based on the high solubility of the chimeras and the autoregulation mechanism of FUS and TDP-43, we propose that these chimeric proteins can be developed as highly soluble and functional replacements for endogenous disease-causing FUS and TDP-43. We demonstrated using i^3^-neurons that FUS_500-IBB_ and FUS_526-IBB_ successfully downregulate endogenous FUS expression level, replace essential FUS functions in DNA-damage repair and RNA processing, and restore neuronal fitness. Similarly, TDP-43_M9M_ restores the disrupted cryptic exon repressing function of TDP-43 in patient-derived fibroblasts harboring mutant TDP-43. Moreover, results from TDP-43_M9M_- and TDP-43_IBB_-expressing cells suggest that these chimeric proteins could promote the nuclear localization of endogenous TDP-43, providing a tool to study the highly disease-relevant mechanism of TDP-43 nuclear transport. Together, our study paves the way for the development of chimeric proteins as therapeutic agents to address the loss- and gain-of-function toxicity in neurodegenerative diseases.

### Autoregulatory feedback enables functional replacement without overexpression toxicity

A striking feature of the chimeric RBPs described here is their ability to reduce endogenous FUS or TDP-43 expression while preserving nuclear function. Both FUS and TDP-43 are known to autoregulate their own levels through binding to nascent transcripts, thereby maintaining tight control over protein abundance^31–36^. Our results indicate that soluble, nuclear chimeric RBPs efficiently engage this autoregulatory circuitry, leading to downregulation of the endogenous, aggregation-prone protein and a concomitant restoration of RBP homeostasis toward a functional replacement.

This mechanism addresses both LoF and GoF toxicity simultaneously and distinguishes chimeric replacement from either simple overexpression or depletion strategies. Overexpression of wild-type FUS or TDP-43 is well documented to exacerbate aggregation and neuronal toxicity^61–63^, reflecting the narrow physiological window within which these RBPs operate. In contrast, expression of chimeric RBPs is well tolerated, likely the result of a combined effect of autoregulatory feedback that limits total RBP abundance and/or highly nuclear localization of the chimera proteins, which together prevent cytoplasmic accumulation beyond physiological thresholds. Future studies comparing the abundance and localization of exogenously expressed chimeric RBPs or WT RBP, relative to the respective endogenous RBP will likely yield important mechanistic insight on this physiological threshold and guide further therapeutic development.

This replacement strategy also offers distinct benefits over depletion strategies. In familial ALS cases driven by mutant FUS (e.g., FUS^P525L^), antisense oligonucleotide-mediated reduction of the mutant protein can be sufficient to alleviate toxicity^64^. However, the majority of ALS cases are sporadic and lack mutations in FUS or TDP-43^9,65–69^. In these settings, pathological mislocalization and aggregation involve wild-type protein, and indiscriminate depletion, particularly of TDP-43, risks exacerbating loss-of-function phenotypes. By preserving nuclear RBP function while reducing aggregation-prone endogenous protein, chimeric replacement may therefore be particularly well suited for sporadic ALS, where restoring homeostasis rather than eliminating protein is likely essential.

Conceptually, this work highlights a shift from protein depletion or supplementation toward homeostatic reconfiguration. By exploiting endogenous autoregulatory feedback, chimeric RBPs avoid the dual pitfalls of loss-of-function toxicity and overexpression-induced aggregation. This principle may extend beyond FUS and TDP-43 to other autoregulated RBPs, suggesting that feedback-aware protein engineering could provide a general strategy to stabilize protein networks that are otherwise vulnerable to pathological imbalance.

### Additional therapeutically relevant features of TDP-43 chimeras

While both FUS and TDP-43 chimeras downregulate endogenous protein and restore nuclear functions, our results reveal an additional property unique to TDP-43 chimeras. In addition to reducing endogenous TDP-43 levels, chimeric TDP-43 proteins promote nuclear localization of the remaining endogenous TDP-43. This effect is not observed with wild-type TDP-43 expression and likely reflects intrinsic features of TDP-43 biology, including its dimerization and oligomerization domains^70^. Through these interactions, chimeric TDP-43 may stabilize nuclear TDP-43 assemblies and counteract cytoplasmic mislocalization of the endogenous protein, even under conditions that favor cytoplasmic mislocalization. By both downregulating excess endogenous TDP-43 and facilitating nuclear retention of the remaining pool, chimeric TDP-43 may restore protein homeostasis and nuclear function more effectively than strategies that act solely on protein abundance. This dual activity is especially relevant in sporadic ALS, where WT TDP-43 pathology is widespread and not driven by mutations.

### Tunable design of chimeric RBPs enables control over autoregulation and functional output

An important insight from this study is that the activity of chimeric RBPs is tunable, rather than binary. For FUS, modifying the length of the residual PY-NLS altered chimeric behavior, with shorter and longer variants exhibiting different kinetics in recruitment to DNA damage site (Figure 3B-D). These findings indicate that partial retention of native sequence features can modulate chimeric activity, providing a means to fine-tune functional output.

Similarly, for TDP-43, distinct nuclear import receptor-NLS pairings produced chimeras with different activities. M9M- and IBB-based designs differed in their ability to promote nuclear localization of both chimeric and endogenous TDP-43 (Figure 6B and D) and in their *in vitro* solubility^25^ (Figure 5E). These differences highlight that NIR-NLS pairing is not interchangeable, and that matching cargo protein properties with appropriate import machinery is critical for optimal function.

Together, these results demonstrate that chimeric RBP behavior can be systematically adjusted by modifying either the core protein or the appended NIR-recognition motif. This tunability opens the door to future optimization strategies, including mutational scanning of key NLS residues or linker regions to further refine autoregulatory strength, nuclear retention, and functional rescue. More broadly, these findings establish chimeric RBPs as a modular platform in which proteostasis, localization, and function can be independently calibrated to meet specific biological or disease-related constraints.

In summary, our work demonstrates that strengthening NIR-NLS coupling through rational protein engineering enables functional replacement of aggregation-prone RBPs by engineered chimeric proteins with enhanced solubility. These chimeric RBPs retain key nuclear functions while engaging endogenous autoregulatory pathways to reduce aggregation-prone endogenous FUS or TDP-43. As a result, engineered RBP chimeras provide a distinct framework for restoring RBP homeostasis while mitigating both gain- and loss-of-function toxicity. More broadly, our findings establish engineered RBP chimeras as a modular and potentially generalizable therapeutic strategy for ALS and related neurodegenerative diseases characterized by RBP mislocalization and aggregation.

## METHODS

### Plasmid construction

#### Mammalian expression plasmids

FLAG-tagged FUS_IBB_ expression plasmids were constructed based on pFUS_IBB_-FLAG^25^ by inserting missing amino acid sequence (amino acids 501 through 526). mClover-FLAG- tagged FUS expression plasmids were constructed by first amplifying FUS_IBB_ sequence from the pFUS_IBB_-FLAG plasmid and inserting into pcDNA5 plasmid under CMV promoter. wtTDP43tdTOMATOHA was a gift from Zuoshang Xu (Addgene plasmid # 28205; http://n2t.net/addgene:28205; RRID: Addgene_28205). M9M- and IBB-tagged TDP-43 expression plasmids were constructed based on wtTDP43tdTOMATOHA by digesting the plasmid with MfeI-HF (R3589S) and KpnI, then inserting synthetic dsDNA containing GS-linker and M9M/IBB sequence using NEBuilder HiFi DNA Assembly Master Mix (New England Biolabs, E2621S). GFP-tagged TDP-43 expression plasmids were constructed by replacing the tdTomato sequence in wtTDP43tdTOMATOHA with the GFP sequence. The K82Q mutation was introduced to the GFP-tagged TDP-43 plasmid by digesting the base plasmids with XhoI (New England Biolabs, R0146S) and AflII (New England Biolabs, R0520S) and inserting a synthetic dsDNA containing K82Q mutation (Integrated DNA Technologies) using NEBuilder HiFi DNA Assembly Master Mix.

#### Lentiviral transfer plasmids

FUS and TDP-43 open reading frames were cloned into pLenti-hSyn transfer plasmid^71^.

#### Bacterial expression plasmids

Construction of M9M-tagged TDP-43 expression plasmid was explained previously^25^. The TDP-43_IBB_ expression plasmid was made by replacing the M9M sequence in TDP-43_M9M_ with the IBB sequence.

All plasmids were verified by sequencing.

### Cell culture

HEK293 (ATCC, CRL-1573) and HEK293T (ATCC, CRL-3216) cells were cultured in Dulbecco’s Modified Eagle’s Medium (DMEM; Corning, 10-013-CV) supplemented with 10% fetal bovine serum (FBS; Cytiva, SH30396.03) and penicillin-streptomycin (Gibco, 15140-122). U2OS cells (ATCC, HTB-96) were cultured in modified McCoy’s 5 A (Gibco, 16600082) supplemented with 10% FBS and penicillin-streptomycin. SH-SY5Y cells (ATCC, CRL-2266) were cultured in adherent conditions in DMEM/F-12 (Gibco, 11320033) supplemented with 10% FBS. Flp-In T-Rex 293 cells (Invitrogen, R78007) were cultured in DMEM supplemented with 10% FBS and either 10 μg/mL zeocin (Gibco, R25001; for parental cell line) or 200 μg/mL hygromycin B (Gibco, 10687010; for chimeric FUS inducible cell line). 15 μg/mL blasticidin was added every other passage in addition to zeocin and hygromycin B.

### iPS and neuronal cell culture

Cortical and motor neuron cultures were generated from iPSCs using the transcription factor-mediated differentiation method pioneered by Michael Ward and Kevin Talbot^45^. These “i^3^” neurons are integrated, inducible, and isogenic. The control cortical and motor neuron lines used in Figure 1 were kind gifts from Drs. Ward and Talbot. When these cells were originally generated, informed consent was obtained prior to tissue donation and generation of pluripotent stem cells, with the tissue collection conforming to the guidelines dictated in the WMA Declaration of Helsinki and the Department of Health and Human Services Belmont Report. For assessment of FUS_IBB_ impacts in FUS mutation bearing cells, additional cell lines Kolf2.1J (control) and FUS^R495X^ (from the INDI project at Jackson Laboratories) as well as FUS^P525L^ and FUS^P525L^ (isogenic revertant) (from Ludo Van den Bosch, KE Leuven) were also converted to be “i^3^” cortical neurons using the same protocol previous described by Ward^45^. Briefly, a cassette containing the neurogenin-2 transcriptional regulator (NGN2) under a doxycycline-inducible promoter was stably integrated into a safe-harbor locus within these lines. Characterization of these lines revealed that the control Kolf2.1J and FUS^P525L^ revertant lines were indistinguishable in differentiation, growth, and neuronal activity, and have therefore been collapsed into a single “control” comparison for this study to make comparisons across mutations more straightforward. The “i3” motor neurons from Ward and Talbot operate under a similar doxycycline inducible cassette, which contains the NGN2 transcription factor, along with Islet-1 (ISL1) and LIM Homeobox 3 (LHX3) to create cells of the motor neuron lineage.

In the iPS stage, cells were maintained in Stemflex Media (Thermofisher) plus 1% penicillin/streptomycin (pen/strep, Hyclone) on matrigel-coated plates (Corning). For differentiation, cells were dissociated into single cells using accutase (Corning), and switched into induction media (DMEM/F12 media with L-glutamine, N2, NEAA, 2 mg/ml doxycycline) which was changed daily for three days. Cells were then dissociated and placed into experimental plating conditions, on poly-L-ornithine coated plates in Maturation Media (BrainPhys media with BDNF, NT-3, and Laminin). Cells were matured and aged longitudinally, with half media changes twice weekly.

### Collection and Culturing of Fibroblasts

The Weinberg ALS Center at Jefferson maintains a biobank of patient-derived fibroblast samples, in accordance with our IRB, which also operates under the guidelines dictated in the WMA Declaration of Helsinki and the Department of Health and Human Services Belmont Report. After obtaining informed consent, our clinician neurologist acquires a 3-mm skin punch biopsy from the inner calf with local anesthesia. Skin biopsies are then sub-dissected and seeded into individual wells of a 6-well plate (Corning) in DMEM, high glucose (Cytiva) with 20% FBS (Thermo Fisher) and 1% penicillin-streptomycin (Cytiva). Dishes were allowed to grow to confluency at 37°C with 5% CO_2_, with media changes twice weekly. Cells were dissociated into single cells using TrpLE (Thermo Fisher) and replated onto 10-cm dishes. At confluency, cells were again dissociated and frozen in liquid nitrogen for storage. Subsequent retrieval of specific patient lines for passing and experimentation was performed. Cells were used at passage 6-7 for experiments described. Two TDP43 mutation carriers TDP43^I383V^ and TDP43^G287S^ and two age- and sex-matched controls were selected to evaluate in this study. See Figure 7A for full demographic information.

### Production of lentiviral vector

The second-generation lentiviral vector was constructed by transfecting HEK293T cells in a T175 flask with 16 μg psPAX2 (a gift from Didier Trono; Addgene plasmid # 12260; http://n2t.net/addgene:12260; RRID:Addgene_12260), 4 μg pMD2.G (a gift from Didier Trono; Addgene plasmid # 12259; http://n2t.net/addgene:12259; RRID:Addgene_12259), and 8 μg transfer plasmid carrying FUS or TDP-43 sequence under the human synapsin promoter. 72-hour post-transfection, the cell culture media containing lentivirus was collected, spun down, and filtered through 0.45 μm low retention PVDF. Viruses were then concentrated using Lenti-X Concentrator (Takara, 631231) according to the manufacturer’s instructions. Small aliquots of viruses were stored frozen at −80°C. The viral titer was determined using RT-qPCR^72^.

### Expression of chimeric proteins in neurons via lentivirus

iPSC-derived cortical and motor neurons were matured on glass bottom plates (immunofluorescence analysis), or 6-well plates (western blotting and RT-qPCR) as described above. Lentivirus was diluted in Maturation Media to achieve the multiplicity of infection (MOI) of 50, which resulted in ∼85% transduction efficiency. Cortical neurons (Day 10) or motor neurons (Day 7) were incubated with the virus overnight, with a full media change the following morning. Cells were then matured for another 4 days before fixation for immunofluorescence imaging analysis or harvest for Western blotting and RT-qPCR. Longitudinal live imaging was also performed from Day 1 to Day 4 post-transduction for cell survival and neurite length analysis.

### Immunofluorescence imaging

Cells were either grown on gelatin-coated coverslips or glass-bottom plates and subjected to various treatments as described in figure legends. Cells were then washed with PBS, fixed with 4% paraformaldehyde, and permeabilized with 0.1% Triton X-100. After washing and blocking in 2% bovine serum albumin (BSA, dissolved in PBS), cells were stained with primary antibody for 16 hours at 4 °C. Cells were then washed with 2% BSA and incubated with secondary antibody for 1 hour at room temperature. Finally, cells were washed with PBS and mounted with Vectashield Antifade Mounting Medium with DAPI (Vector Laboratories, H-1200). The following antibodies and ligands were used for imaging experiments: anti-Map2 (Abcam Cat# ab5392, RRID:AB_2138153, 1:5,000), anti-ChAT (Millipore RRID:AB_2079751, 1:1000), anti-FUS (Bethyl Laboratories, A300-294A, 1:1000), anti-FLAG (Sigma Aldrich, f1804, 1:1000), anti-G3BP1 (Invitrogen, PA5-29455, 1:500), anti-TDP-43 (Proteintech, 10782-2-AP, 1:500), Alexa Fluor488-tagged anti-mouse IgG (Invitrogen, A-21202, 1:2000), Alexa Fluor647-tagged anti-mouse IgG (Invitrogen, A-21235, 1:1000), Alexa Fluor594-tagged anti-rabbit IgG (Invitrogen, A-11012, 1:500), Alexa Fluor647-tagged anti-rabbit IgG (Invitrogen, A-21245, 1:1000), and Janelia Fluor 646 HaloTag Ligand (Promega, HT1060, 200 nM). Fluorescence images were acquired by Leica DMi8 inverted microscope (Leica) with a 40x oil objective, or Nikon A1R confocal microscope with 40x or 60x oil objective. Images were analyzed using Fiji^73^ (v2.15.0) and CellProfiler^74^ (v4.2.6).

### Western blot

Samples were prepared by lysing cells in PBS containing benzonase (Millipore, E1014), Halt protease inhibitor cocktail (Thermo Scientific, 78429), and 1% Triton X-100 for 20 min on ice. 4x Sample Buffer was then added to the lysate, and all samples were boiled before loading onto a gel. Fractionated proteins were transferred onto a PVDF membrane (Millipore, IPFL00010) and blocked in TBST (20 mM Tris-HCl pH 7.4, 150 mM NaCl, and 0.1% Tween 20) containing 3% non-fat dry milk for 30 min. The membrane was incubated with primary antibody overnight at 4 °C, washed, and incubated with secondary antibody for 1 hour at room temperature. Primary and secondary antibodies used are as follows: anti-FUS C-terminus antibody (Bethyl Laboratories, A300-294A, 1:5000), anti-FLAG antibody (Sigma-Aldrich, F1804, 1:2000), anti-β-actin antibody (Cell Signaling, 4970, 1:10,000), anti-GFP antibody (Proteintech, 66002-1-Ig, 1:10,000), anti-HA antibody (Sigma-Aldrich, H9658, 1:10,000), anti-GAPDH antibody (Fitzgerald Industries, 10R-G109a, 1:10,000), anti-mouse IgG antibody (Li-Cor, 926-68070, 1:10,000), and anti-rabbit IgG antibody (Li-Cor, 926-32211, 1:10,000). The membrane was imaged with BioRad ChemiDoc MP Imaging System (BioRad), and band intensities were quantified by ImageJ^73^ (v2.14.0). For uncropped blots, see Source Data.

### Reverse transcription and quantitative PCR (RT-qPCR)

RNAs were purified from cells using TRIzol Reagent (Invitrogen, 15596026) and RNA Clean & Concentrator kit (Zymo Research, R1013). 100∼500 ng purified RNA was used per 10 μL reverse transcription reaction together with Random Primers (Invitrogen, 48190011), Deoxynucleotide Solution Mix (New England BioLabs, N0447S), and SuperScript IV Reverse Transcriptase (Invitrogen, 18090010) according to manufacturer’s instructions. Each qPCR reaction contained 2 μL 10-fold diluted reverse transcription reaction, 500 nM primers, and 1x PowerTrack SYBR Green Master Mix (Applied Biosystems, A46012) in 20 μL total volume. PrimeTime qPCR primers were purchased from Integrated DNA Technologies (βActin, Hs.PT.39a.22214847; FUS, Hs.PT.58.40858500.g). The FUS primer only detects the endogenous FUS since the exogenous FUS carries a codon shuffled sequence on the primer binding region. Other primers used in this study are listed in Supplementary Table. qPCR reaction was run on QuantStudio 3 (Applied Biosystems) according to the manufacturer’s instructions. Ct calculations were done using QuantStudio Design & Analysis Software (v1.5.2; Applied Biosystems).

### Doxycycline inducible mClover-tagged FUS_IBB_ cell line

Flp-In T-Rex 293 cells (Invitrogen, R78007) were plated onto 100-mm dish and transfected with 12.6 μg pOG44 (Invitrogen, V600520) and 1.4 μg pcDNA5 that carries FUS_IBB_ open reading frame, using Lipofectamine 3000 (Invitrogen, L3000008). Two-day post-transfection, cells were selected in DMEM containing 10% FBS and 200 μg/mL hygromycin B. All colonies were pooled and maintained in hygromycin B-and blasticidin-containing medium as explained above.

### Transfection, bortezomib (BTZ), and stress

HEK293, HEK293T, and U2OS cells were transfected using Lipofectamine 3000 Transfection Reagent (Invitrogen, L3000001) according to the manufacturer’s instructions. For immunofluorescence, Western blot, and RT-qPCR experiments, 2.5 μg (6-well) and 1 μg (12-well) plasmid DNA was used. For knockdown experiments, 75 pmol siRNA (Integrated DNA Technologies; control siRNA 51-01-14-04 and siRNA against FUS^32^) was used per 6-well and 35-mm glass bottom dish.

Fibroblasts used for TDP43_M9M_ experiments were also transfected using Lipofectamine 3000 Transfection Reagent according to the manufacturer’s instructions. For immunofluorescence, Western blot, and RT-qPCR experiments, 1.25 μg plasmid DNA was used per 6-well.

For proteasomal inhibition experiments, HEK293T cells were treated with 200 nM BTZ (or equal volume DMSO) for 24 hours. The BTZ treatment and transfection were done simultaneously where appropriate.

To induce osmotic stress, the growth media was replaced with 0.4 M sorbitol containing media, and cells were incubated for 1 hour at 37°C and 5% CO2. To induce oxidative stress, the growth media was replaced with 500 μM sodium arsenite containing media, and cells were incubated for 1 hour at 37°C and 5% CO2.

### Microirradiation assay

The microirradiation assay was conducted according to the previously published protocol with minor modifications. The doxycycline-inducible HEK293 cells carrying FUS cassette were plated on 35-mm glass bottom dish and treated with 100 ng/mL doxycycline and siRNA for 48 hours. 30 minutes before imaging, cells were synthetized to DNA damage by addition of 2 μM Hoechst 33342 (Thermo Scientific 62249). Irradiation and imaging were conducted using a Nikon A1R confocal microscope with a resonant scanner, 40x oil objective, and 405-nm laser at 10% power. Images were acquired continuously for 180 seconds, and mean fluorescence intensity per pixel was calculated within regions of interest using Fiji^73^.

### DNA repair homologous recombination (HR) reporter assay

HR efficiency was assessed using DR-U2OS cells containing a single integrated DR-GFP reporter^48,49^. Exponentially growing cells were seeded in 6-well plates at a density of 2 × 10^5^ cells per well and reverse-transfected with 2 μl siRNA and 3 μl Lipofectamine™ RNAiMAX (Invitrogen, 13778150). After 24 hours, cells were transfected with 1.5 μg of the indicated expression plasmids using 3 μl Lipofectamine™ 2000 (Invitrogen, 11668027). On the following day, cells were transfected with 1 μg of the I-SceI endonuclease vector (pCBASce) using 2 μl Lipofectamine™ 2000. HR-mediated repair of the I-SceI-induced double-strand break restored GFP expression, which was quantified by flow cytometry (BD FACSCalibur) 72 hours after I-SceI transfection. HR efficiency was calculated as the percentage of GFP-positive cells relative to control conditions.

### COMET Assay

COMET assays to assess levels of DNA damage in cortical i^3^ neurons were performed according to manufacturer’s instructions with alkaline electrophoresis (OxiSelect Comet Assay, Cell Biolabs Inc). Briefly, cortical i^3^ neurons were transduced on Day 10 with GFP or GFP-tagged FUS_IBB_, as described above, and were harvested on Day 14 for DNA damage assessment. Cells were removed by accutase (Corning) dissociation from experimental plating conditions and were resuspended at a concentration of 1 x 10^5^ cells/mL in ice-cold PBS. These cells were then combined with the COMET Agarose at a 1:10 ratio (v/v), with 75 μL of this mixture added to the comet slides. Slides were incubated in the dark at 4°C for 30 minutes, and then immersed in lysis buffer for 60 minutes again in darkness at 4°C. Following lysis, slides were switched into pre-chilled alkaline solution for 30 minutes, in darkness, at 4°C. Slides were then placed into a horizontal electrophoresis apparatus filled with alkaline solution, and were run for 20 minutes at 1volt/cm. Following three 2-minute pre-chilled deionized H_2_O washes, slides were placed in pre-chilled 70% ethanol for 5 minutes. Slides were then incubated at 37°C for 30 minutes to dry. Once totally dry, 100 μL/slide well of Vista Green DNA Dye was added and incubated for 15 minutes. Slides were then visualized with the Nikon Eclipse Ti microscope. Images were evaluated for Comet Phenotypes using Fiji^73^ and the “Open Comet” plugin^75^. Data is represented as percent tail DNA.

### RNA-seq

FUS and GFP expressions were transduced via lentivirus (MOI = 50) in iPSC-derived cortical neurons at day 10 post-differentiation as described earlier. After 4 days of expression, cells were lysed in TRIzol Reagent and submitted for library preparation and sequencing at Novogene. Strand-specific libraries were prepared from Oligo(dT) enriched samples, and sequencing reactions were carried out on Illumina NovaSeq X Plus platform to produce >60 million paired-end 150 reads per sample. Cleaned reads were mapped to GRCh38 primary assembly using HISAT2^76^, which resulted in >50 million mapped reads per sample. Mapped reads were further processed by featureCounts^77^ and DESeq2^78^ for differential expression analyses. Three biological replicate samples were sequenced per treatment condition.

### Longitudinal live-cell imaging survival analysis (Viability Assay)

Control cortical i^3^ neurons were transduced with GFP-tagged FUS (Sup. Figure 4) on DIV 10 at an MOI= 50 and underwent longitudinal live-cell imaging for two weeks to assess survival and risk of death. This assay was performed as previously described^79,80^, using a Nikon Eclipse Ti microscope and CoolSNAP ES2 high-performance CCD camera, with all aspects of stage/shutter movements and imaging acquisition controlled by NIS-Elements Imaging software (Nikon). GFP fluorescence was used as a marker of cell transduction, and DRAQ7 dye (Abcam, 3 μM) added to the media indicates loss of membrane integrity and cell death. Visual inspection of image files generated from the automated protocol allows for evaluation of individual neurons over time, with death defined as the day on which the neuron picked up DRAQ7 positivity.

### Assessment of neurite length and complexity

Cortical i^3^ neurons were transduced with either GFP-tagged FUS or tdTomato-tagged TDP43 on Day 10 at an MOI = 50 and were evaluated following 4 days of transgene expression. Immunostaining with neuronal marker anti-Map2 was performed as described above, with AlexaFluor-546 (for GFP-tagged FUS containing cells) or AlexaFluor-488 (for tdTomato-tagged TDP43 containing cells) secondary antibodies. Following imaging by confocal microscopy, neuron branch tracing was performed using Fiji^73^ equipped with the Neuroanatomy SNT plugin^81^ to determine total branch length.

### Purification of MBP tag and TDP-43-MBP from bacterial culture

TDP-43 protein expression and purification were described previously^25,82,83^. Briefly, BL21-CodonPlus (DE3)-RIL E. coli cells were transformed with TDP-43 expression constructs, and the protein expression was induced with 1 mM IPTG for 18 h at 16°C. Cells were pelleted, resuspended in TDP-43 Lysis Buffer (20 mM Tris-HCl pH 8.0, 1 M NaCl, 10 mM imidazole, 10% glycerol, 1 mM DTT, and cOmplete EDTA-free Protease Inhibitor Cocktail), and lysed by sonication. The lysate was clarified by centrifugation and bound to HisPur Ni-NTA Resin. The crude protein was eluted with TDP-43 Elution Buffer (20 mM Tris-HCl pH 8.0, 1 M NaCl, 500 mM imidazole, 10% glycerol, and 1 mM DTT). The protein was further purified by size exclusion chromatography using Superdex 200 increase 10/300 GL column (GE Healthcare) with TDP-43 SEC Buffer (20 mM Tris-HCl pH 8.0, 300 mM NaCl, and 1 mM DTT). A typical A260/A280 ratio for the purified protein was around 0.6. The free MBP protein was purified similarly from the same vector without TDP-43 sequence.

### Importin β1 purification from bacterial culture

Importin β1 was purified from the importin α1/β1 dimeric complex^70^. Briefly, pACYCDuet-1 plasmids encoding importin α1/β1 sequences were transformed into BL21-DE3 E. coli. The importin α1/β1 dimeric complex were expressed for 3 h at 28 °C with 500 µM IPTG, lysed in Imp α/β Lysis Buffer (20 mM Tris-HCl pH 8, 75 mM NaCl, 3 mM β-mercaptoethanol, and 0.1 mM PMSF), and bound to Low Density Nickel Agarose (Gold Biotechnology, H-321-100). The complex was washed using Imp α/β Low Salt Buffer (20 mM Tris-HCl pH 8, 50 mM NaCl, 3 mM β-mercaptoethanol, and 0.1 mM PMSF), and Impβ1 was eluted by using Imp α/β High Salt Elution Buffer (20 mM Tris-HCl pH 8, 250 mM MgCl2 and 150 mM NaCl). The protein was further purified by size-exclusion chromatography. A typical A260/A280 ratio for the purified protein was around 0.6.

### Kapβ2 purification from bacterial culture

His-SUMO-Kapβ2 was expressed and purified as previously described^84^. The expression of protein in BL21-CodonPlus (DE3)-RIL cells was induced by 1 mM IPTG for 16 hours at 16°C. The cell pellet was resuspended and lysed in Resuspension Buffer (50 mM Tris-HCl pH 7.5, 100 mM NaCl, 20% glycerol, 10 mM imidazole, 2.5 mM β-mercaptoethanol, and cOmplete EDTA-free Protease Inhibitor Cocktail). The lysate was clarified by centrifugation, incubated with HisPur Ni-NTA Resin, washed, then eluted with Kapβ2 Elution Buffer (50 mM Tris-HCl pH 7.5, 100 mM NaCl, 20% glycerol, 200 mM imidazole, and 2.5 mM β-mercaptoethanol). Next, His-SUMO tag was cleaved by incubating with ULP1, then Κapβ2 was further purified by HiTrap Q HP column (GE Healthcare, 17-1154-01) with gradient of Kapβ2 Buffer A (20 mM imidazole pH 6.5, 75 mM NaCl, 20% glycerol, and 2 mM DTT) and Kapβ2 Buffer B (20 mM imidazole pH 6.5, 1 M NaCl, 20% glycerol, and 2 mM DTT). A typical A260/A280 ratio for the purified protein was around 0.6.

### TDP-43 inhibition assay

The inhibition assay for MBP-tagged TDP-43 has been described previously^85^. MBP-tagged TDP-43 protein was thawed on ice and buffer exchanged into TDP-43 Turbidity Assay Buffer (20 mM HEPES-NaOH pH 7.4, 150 mM NaCl, and 1 mM DTT) using Amicon Ultra Centrifugal Filter, 30 kDa MWCO (Millipore, UFC5030). The buffer exchanged protein was then centrifuged for 10 min at 23,000 × g, 4 °C. Clear supernatant was transferred to a clean tube, and the protein concentration was estimated by Bradford assay. At time = 0 min, 5 μM TDP-43, 32 μg/mL TEV, and importins (where appropriate) were mixed in TDP-43 Turbidity Assay Buffer. The total reaction volume was 100 μL, and all reactions were carried in a clear-bottom, black-well 96-well plate. OD395 measurements were taken by TECAN Spark plate reader.

### TDP-43 disaggregation assay

MBP-tagged TDP-43 protein was thawed on ice and buffer exchanged into TDP-43 Turbidity Assay Buffer using Amicon Ultra Centrifugal Filter, 30 kDa MWCO. The buffer exchanged protein was then centrifuged for 10 min at 23,000 × g, 4°C. Clear supernatant was transferred to a clean tube, and the protein concentration was estimated by Bradford assay. At time = 0 min, 5 μM TDP-43 and 16 μg/mL (WT and TDP-43_IBB_) or 32 μg/mL TEV (TDP-43_M9M_) were mixed in TDP-43 Turbidity Assay Buffer. The different amount of TEV accounts for the differences in cleavage efficiency^25^. The total reaction volume was 90 μL at time = 0 min, and all reactions were carried in a clear-bottom, black-well 96-well plate. OD395 measurements were taken by TECAN Spark plate reader. At time = 120 min, 10 μL Kapβ2 or matching buffer was added to each reaction and mixed gently by pipetting up and down. The turbidity measurements were resumed immediately and recorded for 120 more minutes.

### TDP-43 anisotropy assay

MBP-tagged TDP-43 was diluted in Anisotropy Assay Buffer (20 mM HEPES-NaOH pH 7.4, 150 mM NaCl, and 1 mM DTT), then fluorescein-tagged RNA (synthesized by Horizon Discovery) was added to final concentration of 10 nM. The protein-RNA mixture was incubated 20 min, and fluorescence anisotropy was measured at 25 °C by TECAN Spark plate reader. Triplicate measurements were then plotted and fitted as previously described^25^.

### Statistical analysis

All statistical analyses were carried out by GraphPad Prism, and methods used are indicated in figure legends.

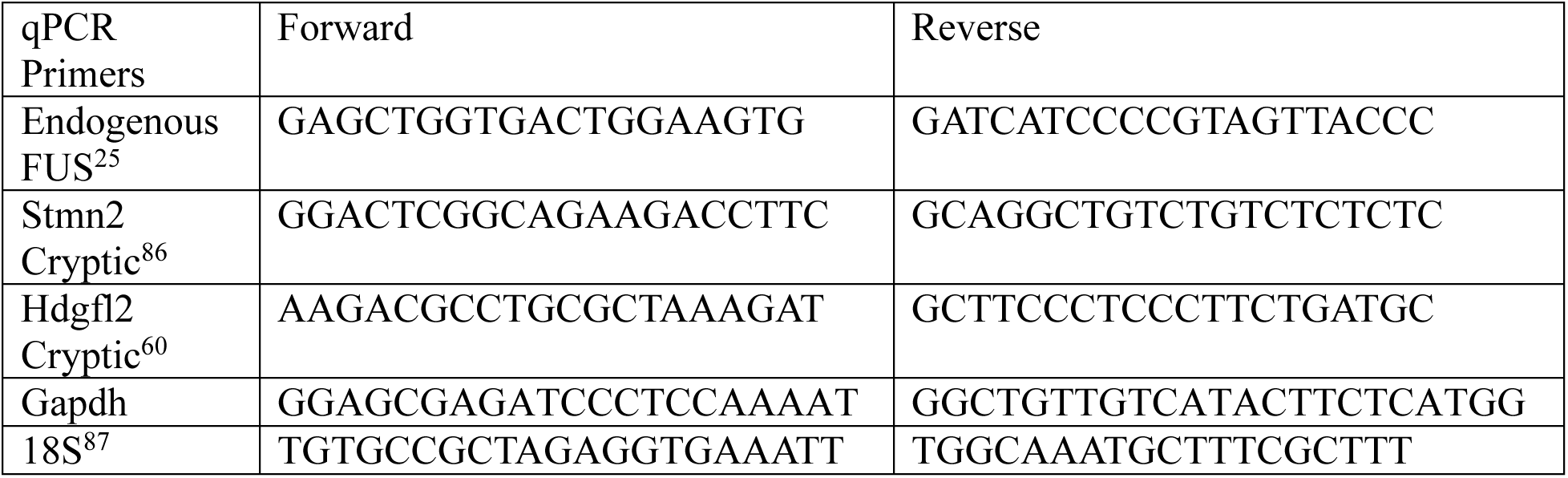

## ACKNOWLEDGEMENT

We thank Dr. Sami Barmada for kindly sharing the HEK293T cell line carrying Halo-tagged TDP-43. We also thank Drs. Jason Hill and James Muller at the BioImaging Shared Resources of the Sidney Kimmel Comprehensive Cancer Center (NCI 5 P30 CA-56036) at Thomas Jefferson University for their assistance with live-cell imaging experiments. L.G. is supported by the National Institute of General Medical Sciences grant R35GM138109, and the National Institute of Neurological Disorders and Stroke grant R01NS121143. B.K.J. and P.P. are supported by The Farber Family Foundation and Family Strong 4 ALS. W.Z. is supported by R01GM141091, American cancer society research scholar grant RSG-22-721675-01-DMC, and discovery boost grant DBG-25-1292405-01-DMC.

## AUTHOR CONTRIBUTIONS

M.H., B.K.J., T.N., and W.L. performed experiments and analyzed data. A.G., S.S.M., Y.K., N.A., and J.A.D. performed experiments. M.H., B.K.J., and L.G. wrote the manuscript. W.Z., G.C., P.P., and L.G. supervised the work.

## MATERIALS & CORRESPONDENCE

Correspondence and material requests should be made to Piera Pasinelli or Lin Guo.

