## Supplementary Figures for "Engineered Chimeric Proteins Functionally Replace ALS-causing Endogenous FUS and TDP-43 to Improve Neuronal Health"

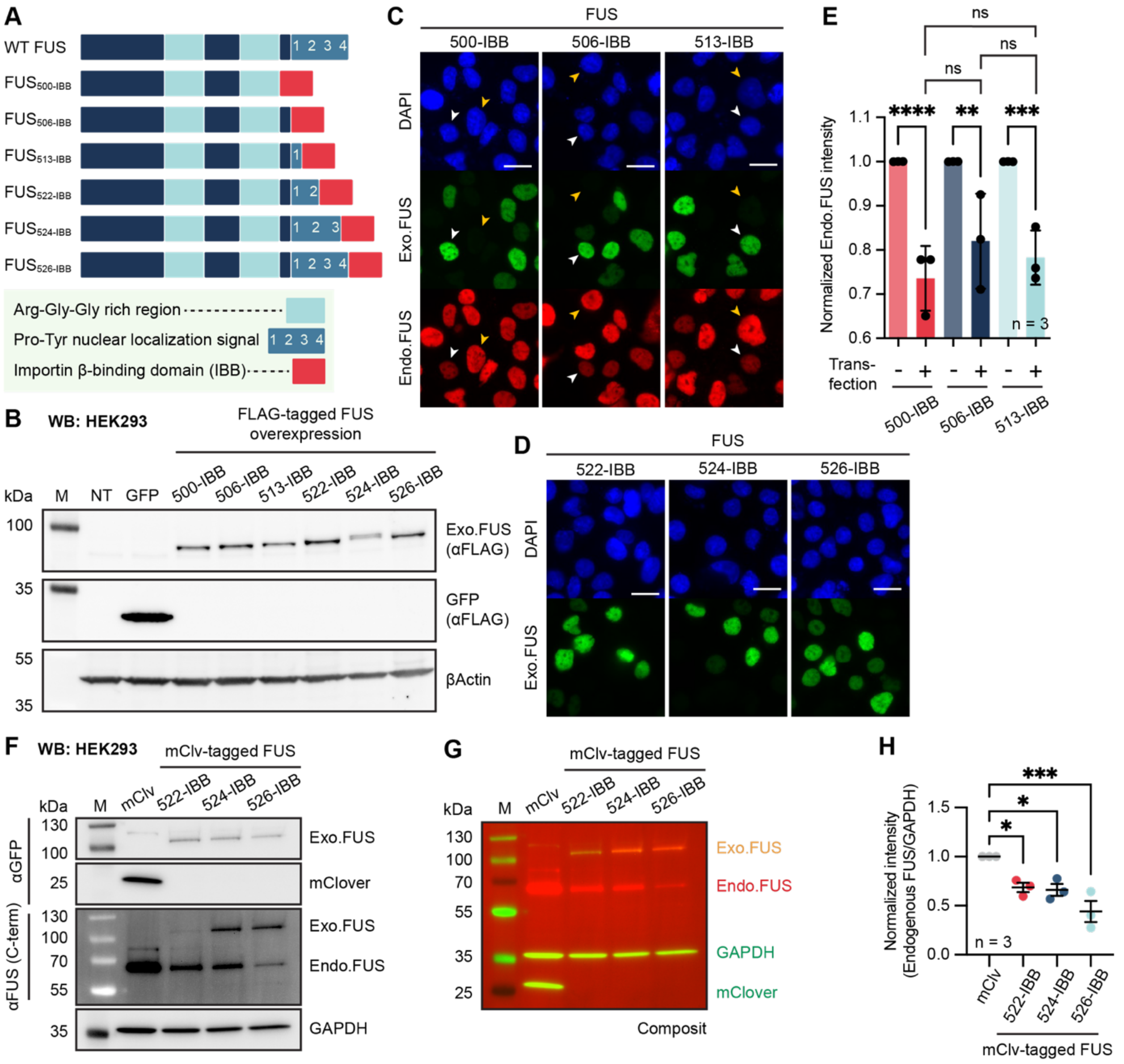


**Supplementary Figure 1: Various FUS-IBB chimeras downregulate endogenous FUS levels**.
**A.** Schematic of FUS domain structure highlighting the different PY-NLS truncations used in the following experiments. The numbers within the PY-NLS box indicate the different epitopes described previously[1].
**B.** Representative Western blot of lysates from HEK293 cells transfected with the indicated FLAG-tagged FUS-IBB constructs. The experiments were repeated three times with similar results. M = molecular weight marker and NT = non-transfected.
**C.** Representative immunofluorescence images of HEK293 cells transfected with the indicated FLAG-tagged FUS-IBB constructs. Cells were fixed and stained with anti-FLAG and anti-FUS PY-NLS antibody. White arrowheads represent transfected nuclei, and yellow arrowheads indicate non-transfected nuclei. Scale bars = 20 µm.
**D.** Representative immunofluorescence images of HEK293 cells transfected with the indicated FLAG-tagged $FUS_{IBB}$ constructs. Cells were fixed and stained with anti-FLAG antibody. The experiment was repeated at least three times with similar results. Scale bars = 20 µm.

**E.** Quantification of images collected in C showing the endogenous FUS fluorescence intensity. Each point represents an independent experiment and is normalized to the non-transfected cells. ** $p < 0.01$, *** $p < 0.001$, **** $p < 0.0001$, and ns = not significant by One-way ANOVA Sidak's multiple comparisons test.

**F.** Representative Western blot of lysates from HEK293 cells transfected with the indicated mClover- and FLAG-tagged FUS-IBB constructs. Exo. FUS = exogenous FUS tagged with mClover and Endo. FUS = endogenous FUS.

**G.** Composite representative Western blot of HEK293 lysates shown in F. Yellow color of the exogenous FUS (Exo. FUS) indicates the overlapping signals from the green (anti-mClover/GFP antibody) and the red (anti-FUS antibody) channels.

**H.** Quantification of the endogenous FUS bands shown in F and G. Mean and SEM of $n = 3$ experiments are shown. Each point represents an independent experiment. * $p < 0.05$ and *** $p < 0.001$ by one way ANOVA Dunnett's multiple comparisons test.

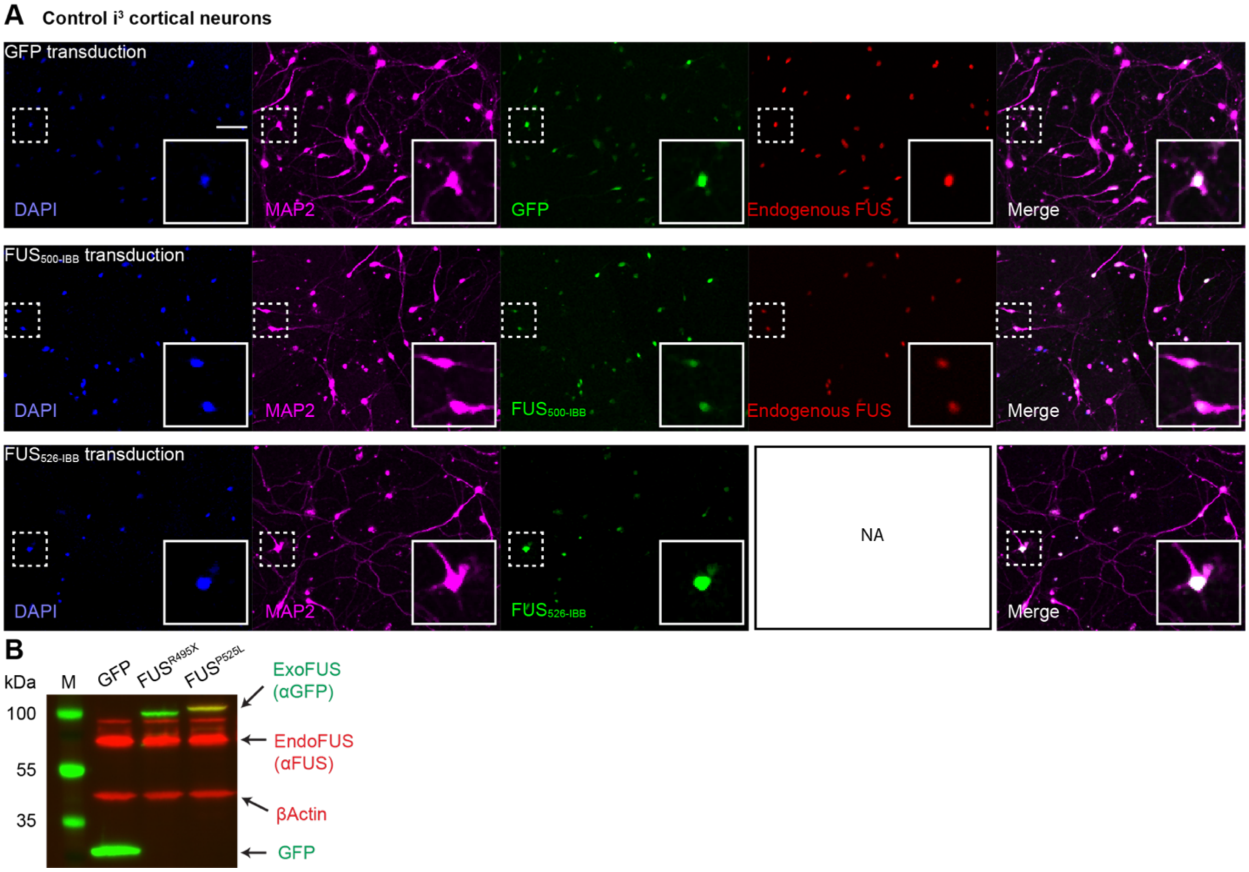


**Supplemental Figure 2: FUS$_{500\text{-IBB}}$ and FUS$_{526\text{-IBB}}$ downregulate endogenous FUS expression in i$^3$ cortical neurons carrying FUS$^{R495X}$ and FUS$^{P525L}$ mutation.**

**A.** Representative images of control i$^3$-CN (Kolf 2.1J) transduced with GFP, GFP-FUS$_{500\text{-IBB}}$, or GFP-FUS$_{526\text{-IBB}}$. Note that the endogenous FUS signal is not shown in FUS$_{526\text{-IBB}}$ transduced cells (denoted as NA) because the anti-FUS antibody detects both FUS$_{526\text{-IBB}}$ and endogenous FUS. Scale bars = 50 μm.

**B.** Representative Western blot of lysates from HEK293 cells transfected with GFP, GFP-tagged FUS$^{R495X,}$ or GFP-tagged FUS$^{P525L}$. FUS$^{R495X}$ does not react with the anti-FUS antibody, whereas FUS$^{P525L}$ shows an overlapping signal (yellow) from anti-FUS (red) and anti-GFP (green) antibody.

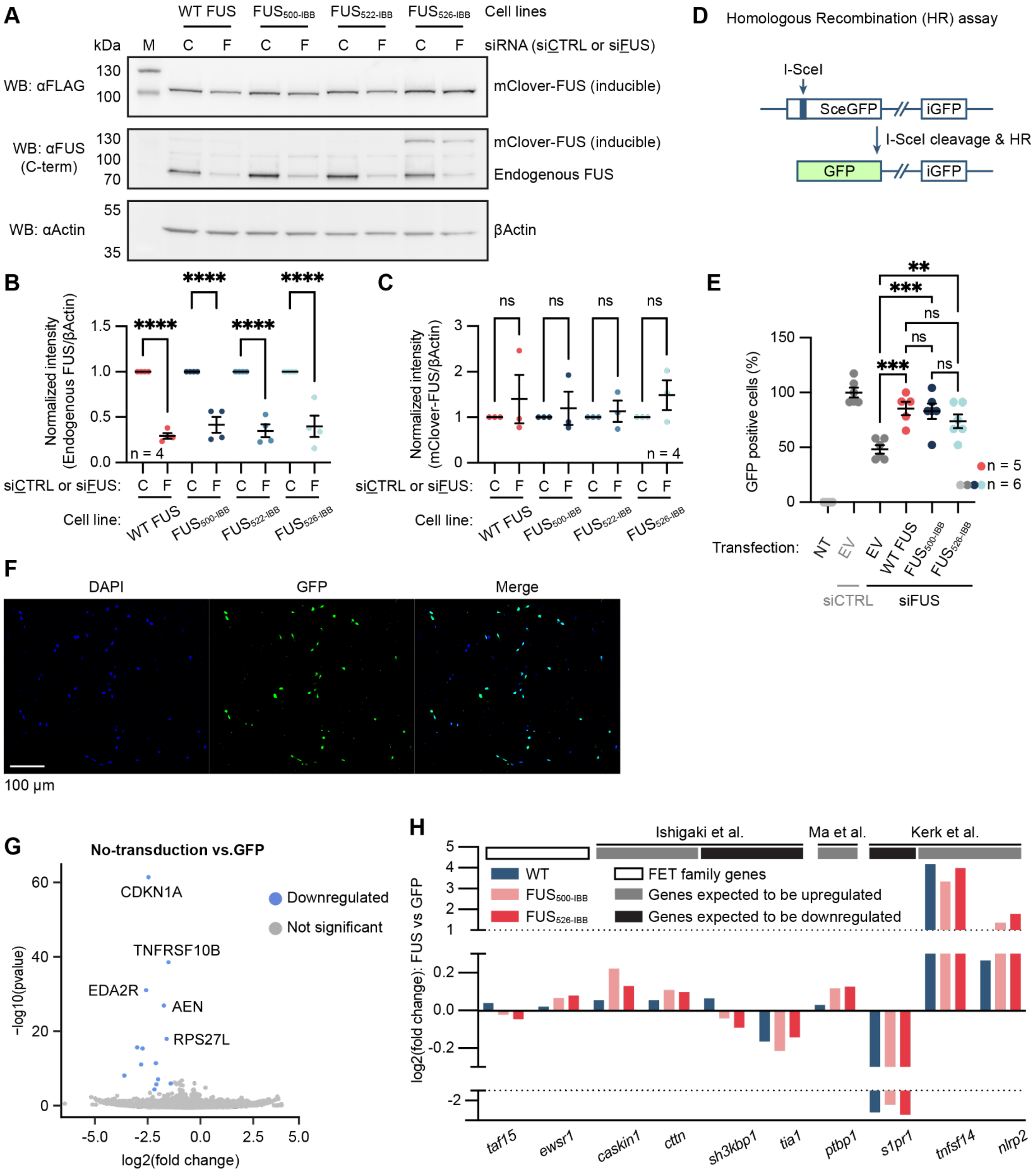


**Supplementary Figure 3: FUS$_{500\text{-IBB}}$ and FUS$_{526\text{-IBB}}$ participate in DNA repair and regulate mRNA expression similarly to WT FUS.**

**A.** Representative Western blot of HEK293 cells expressing the indicated FUS chimera with mClover tag. mClover-FUS expression was induced by the addition of doxycycline to the cell culture media. Cells were treated with either control siRNA (“C”) or FUS siRNA (“F”).

**B and C.** Quantification of A showing the intensity of the endogenous FUS bands (B) or the mClover-tagged FUS chimera band (C) of the indicated cell lines treated with the indicated siRNA. Mean and SEM of four independent experiments are plotted. ns = not significant and **** $p < 0.0001$ by one-way ANOVA Sidak’s multiple comparisons test.

**D.** Design of the homologous recombination (HR) reporter assay. Cleavage of the reporter GFP by I-SceI endonuclease triggers the repair of the GFP open reading frame using the downstream promoter-less template (iGFP) via HR. Successful repair results in GFP protein production.

**E.** Percentage of GFP-positive cells in the indicated HR reporter assay. Cells were treated with either control siRNA (siCTRL) or siRNA against FUS (siFUS) and transfected with empty vector (EV), WT FUS, $FUS_{500\text{-}IBB}$, or $FUS_{526\text{-}IBB}$. Mean and SEM of two independent experiments each containing two or three transfection replicates are plotted. NT = no treatment (no I-SceI, siRNA, or FUS transfection). ns = not significant, *** $p < 0.001$, and **** $p < 0.0001$ by one-way ANOVA Sidak's multiple comparisons test.

**F.** Representative immunofluorescence images of GFP transduced $i^3$-CN. Transduction efficiency for the cells was over 85% consistently across experiments. Scale bars = 100 µm.

**G.** Differential expression analysis of GFP transduced compared to non-transduced neurons. Data points are colored based on $\log_2$(fold change) > 1 and significance ($p < 0.05$).

**H.** Differential expression profile of FET family genes and genes known to be regulated by FUS[2-4].

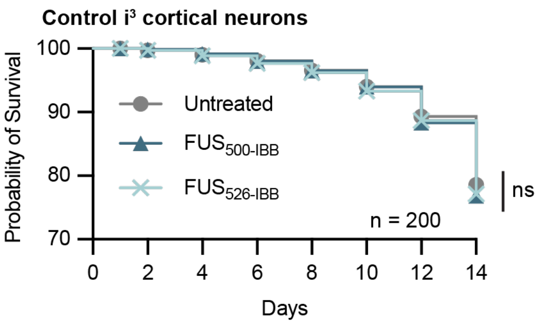


**Supplementary Figure 4: Chimeric FUS expression is not toxic to neurons.**

Longitudinal survival analysis of $i^3$-CN with/without the expression of the indicated chimeric FUS. n = 100 individual neurons were assessed across two independent experiments. ns = not significant by Mantel-Cox test.

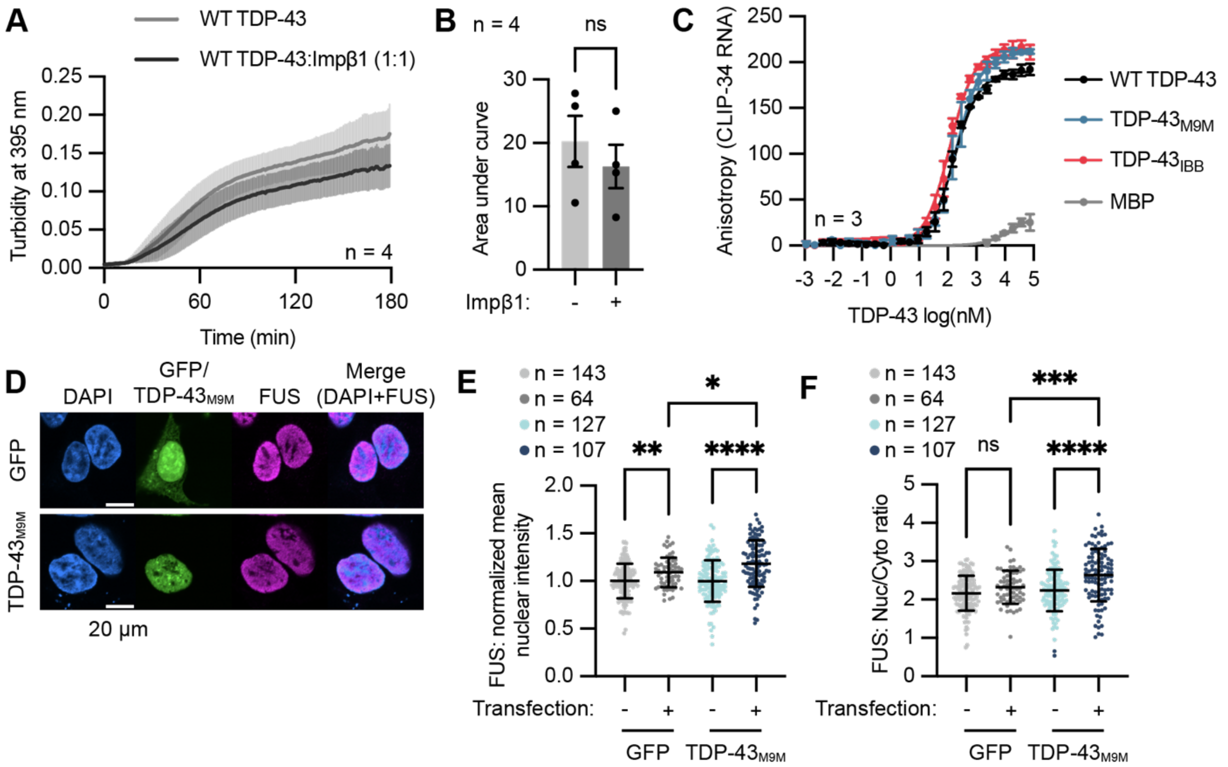


**Supplementary Figure 5: TDP-43$_{M9M}$ can form nuclear condensates and does not disrupt native cargo transport.**

**A.** Turbidity assay curve representing the aggregation profile of WT TDP-43 with and without an equimolar amount of Impβ1. WT TDP-43 and Impβ1 were mixed at the beginning of the reaction when TEV was added to initiate TDP-43 assembly. Turbidity was recorded over time. Mean and SEM of four independent experiments are plotted.

**B.** Quantification of the area under the curve shown in A. Each point represents an independent experiment. ns = not significant by Welch's t test.

**C.** The anisotropy assay with TDP-43 variants. Fluorescein-labeled short RNA, CLIP-34, was added to various concentrations of TDP-43. Each point represents the mean ± SEM from three independent experiments. Fitted curves, which were used to estimate Kd in Figure 5C, are shown as solid lines.

**D.** Representative immunofluorescence images of HEK293 transfected with GFP or GFP-TDP-43$_{M9M}$. Cells were fixed and stained with anti-FUS antibody. Scale bars = 20 µm.

**E.** Quantification of FUS nuclear intensity from the images shown in D. Each point represents a cell and is normalized to the mean of non-transfected cells. * $p < 0.05$, ** $p < 0.01$, and **** $p < 0.0001$ by one-way ANOVA Sidak's multiple comparisons test.

**F.** Quantification of FUS nuclear-to-cytoplasmic (Nuc/Cyto) ratio from the images shown in D. Each point represents a cell. *** $p < 0.001$, **** $p < 0.0001$, and ns = not significant by one-way ANOVA Sidak's multiple comparisons test.

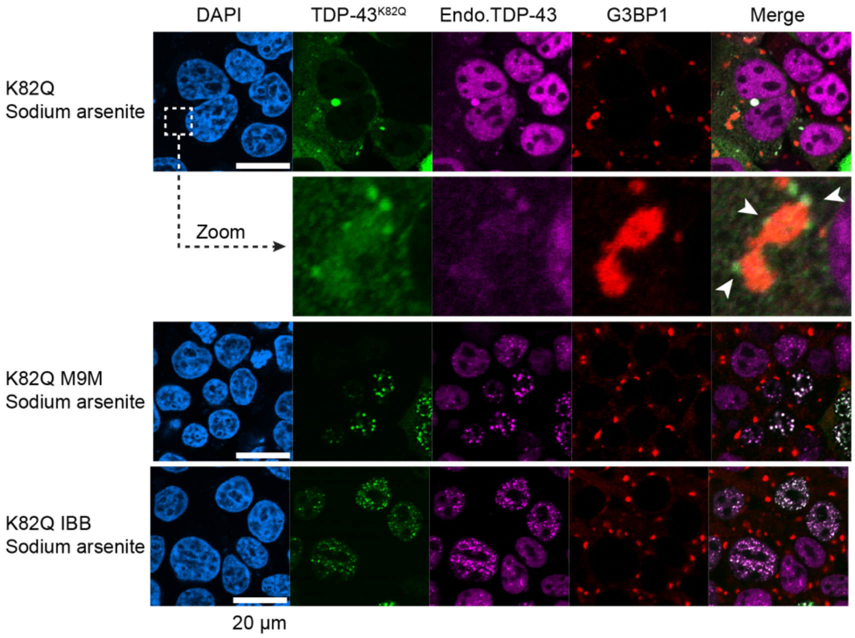


**Supplementary Figure 6: Acetyl-mimetic TDP-43 carrying M9M and IBB localize to the nucleus and can form nuclear condensates.**

Representative immunofluorescence images of HEK293T cells carrying endogenously Halo-tagged TDP-43 and expressing TDP-43 K82Q variants. Cells were treated with sodium arsenite for 1 hour before fixation. White arrowheads indicate TDP-43$^{K82Q}$ puncta adjacent to stress granules. G3BP1 = stress granule marker. Scale bars = 20 µm.

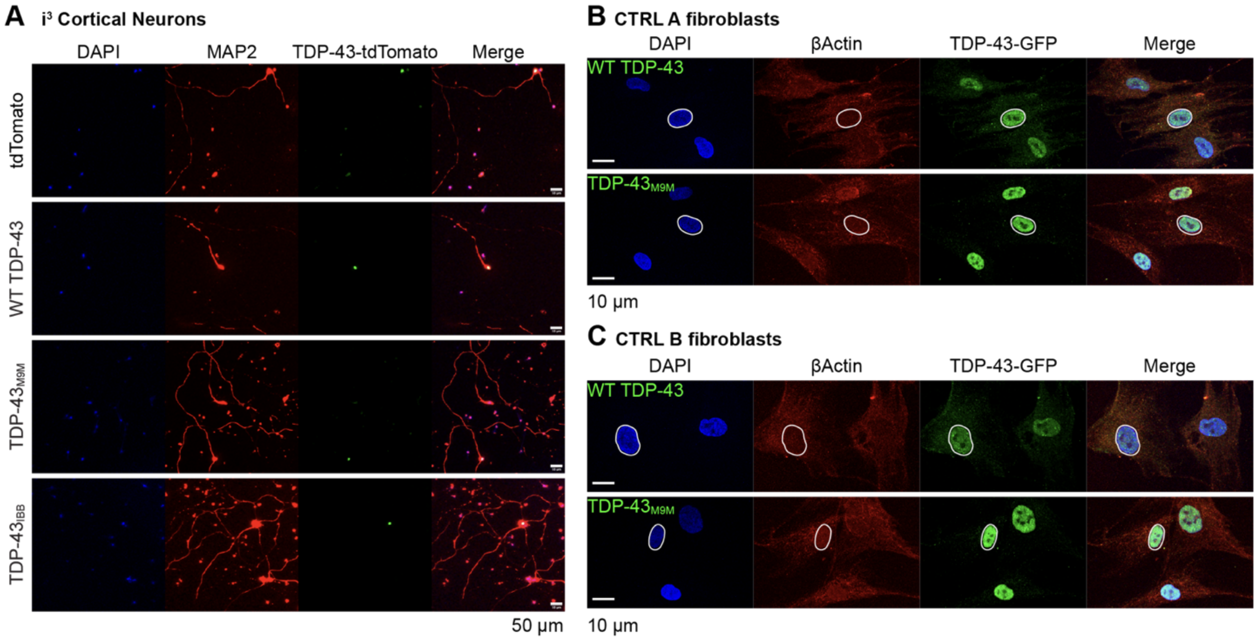


**Supplementary Figure 7: Chimeric TDP-43 localizes in the nucleus of fibroblasts and is not toxic to cortical neurons.**

**A.** Representative immunofluorescence images of control i3-CN transduced with tdTomato-tagged WT or chimeric TDP-43. Images are identical to the main Figure 7A, but this panel includes unmerged channels. Scale bars = 50 µm.

**B and C.** Representative immunofluorescence images of the control patient-derived fibroblasts transfected with GFP-tagged WT or chimeric TDP-43. The nucleus of a representative cell is circled with a white line. Scale bars = 10 µm.
